# Selecting better biocatalysts by complementing recoded bacteria

**DOI:** 10.1101/2022.08.16.504095

**Authors:** Rudy Rubini, Suzanne C. Jansen, Houdijn Beekhuis, Henriëtte J. Rozeboom, Clemens Mayer

**Affiliations:** Biomolecular Chemistry & Catalysis, Stratingh Institute, University of Groningen, Nijenborgh 4, 9747 AG Groningen, The Netherlands; Biotransformation and Biocatalysis, Groningen Biomolecular Sciences and Biotechnology Institute, University of Groningen, Nijenborgh 4, 9747 AG Groningen, The Netherlands

## Abstract

In vivo selections are powerful tools for the directed evolution of enzymes. However, the need to link enzymatic activity to cellular survival makes selections for enzymes that do not fulfill a metabolic function challenging. Here, we present an in vivo selection strategy that leverages recoded organisms addicted to non-canonical amino acids (ncAAs) to evolve biocatalysts that can provide these building blocks from synthetic precursors. We exemplify our platform by engineering carbamoylases that display catalytic efficiencies more than five orders of magnitude higher than those observed for the wild-type enzyme for ncAA-precursors. As growth rates of bacteria under selective conditions correlate with enzymatic activity, we were able to elicit improved variants from populations by performing serial passaging. By requiring minimal human intervention and no specialized equipment, we surmise that our strategy will become a versatile tool for the in vivo directed evolution of diverse biocatalysts.

## Introduction

Enzymes are marvelous catalysts that accelerate reactions with unmatched rates and selectivities.^1,2^ Their use as standalone catalysts or integration into designer microbes promises the sustainable production of fine chemicals, biofuels, and biomedicines.^3,4^ However, enzymes found in nature are rarely optimal for industrial applications with low stabilities, narrow reaction scopes and/or low activities on industrially-relevant substrates restricting their applicability. Courtesy of advances in molecular and structural biology, it has become feasible to tailor enzyme properties by mimicking the Darwinian algorithm in the laboratory (**Fig. 1A**).^5^ One round of the *directed evolution* of biocatalysts comprises three steps: (1) creating diversity by introducing mutations, (2) identifying enzyme variants with improved characteristics and (3) amplifying selected variants. When performed iteratively, these steps enable the gradual improvement of an enzyme’s catalytic performance.6-8 Applying this algorithm for enzyme engineering has proven useful for increasing stability, identifying specialized biocatalysts for non-natural substrates, boosting promiscuous or new-to-nature activities, and altering the stereo- or regioselectivities of enzymes.

**Figure 1:**
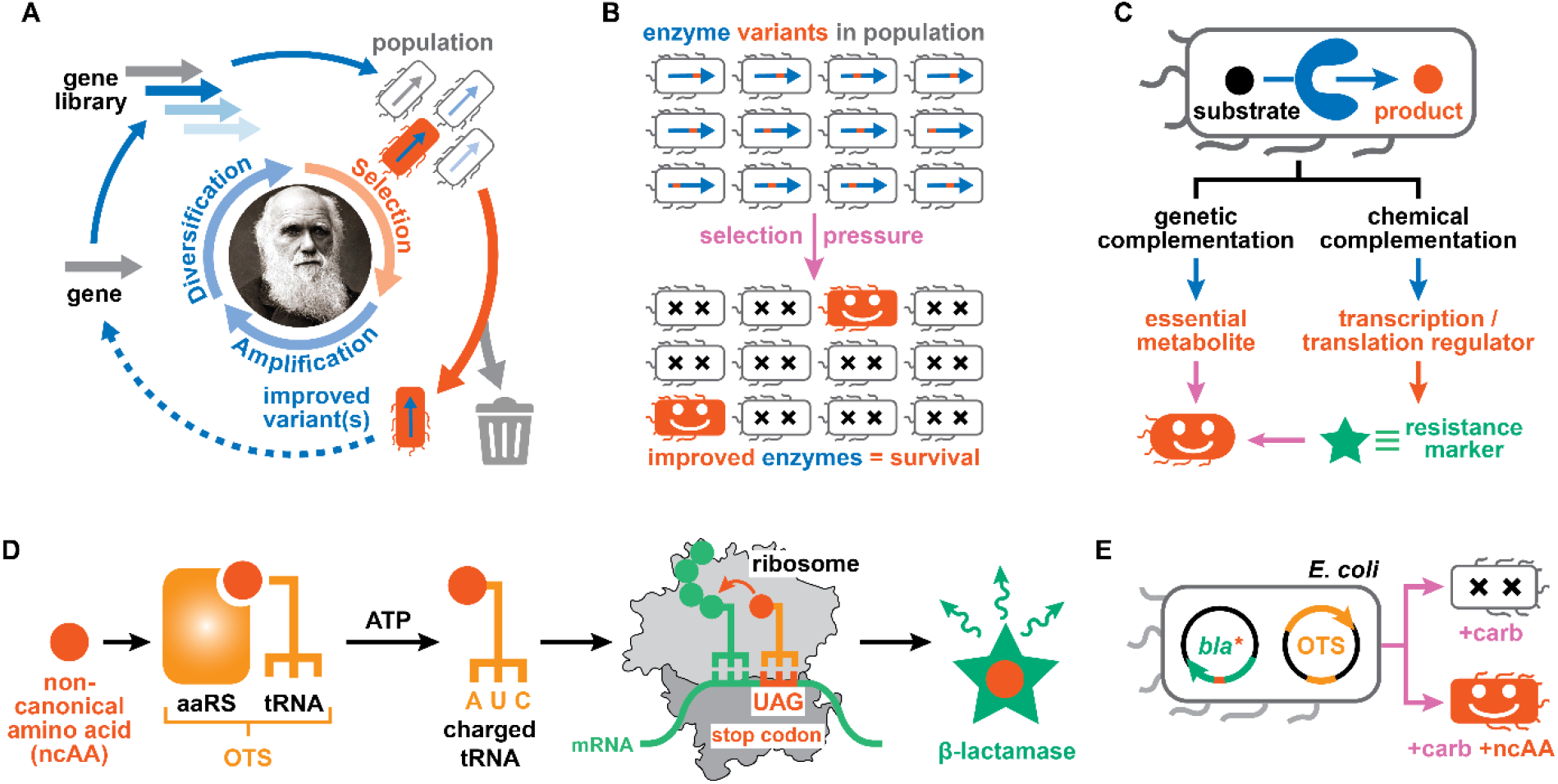
Selection strategies in the directed evolution of biocatalysts. **A:** The properties of biocatalysts are tailored in directed evolution via iterative cycles of gene diversification, selection, and amplification. **B:** In selections, enzyme variants in populations are assessed all at once with an applied selection pressure eliminating all undesired library members. **C:** Selections make use of genetic or chemical complementation strategies to link the activity of an enzyme of interest to the survival/growth of a producing organism. **D-E:** Orthogonal translation systems (OTSs) consist of aminoacyl tRNA synthetase (aaRS) / suppressor tRNA pairs that facilitate the site-selective incorporation of non-canonical amino acids into proteins of interest. When engineering OTSs for ncAAs, functional aaRS/tRNA pairs are identified by allowing *E. coli* to survive in presence of β-lactam antibiotics (e.g. carbenicillin, carb).

Over the past decades, mutagenesis, recombination, and computational tools have sufficiently matured to enable the generation of large gene libraries with billions of variants.^9^ However, navigating the resulting sequence space efficiently remains critically dependent on identifying improved variants based on the (high-throughput) analysis of target products.10 Fundamentally, the strategies employed to analyze gene libraries can be divided into either screens or selections.^6-8^ Screens necessitate the individual assessment of every library member, with throughput thus being limited by the speed of the employed biochemical or biophysical readout.^11,12^ In contrast, selections link the function of an enzyme of interest to the survival or growth of a host organism.^13^ Large populations of up to 10^9^ individual cells can be assessed all at once, as an applied selection pressure eliminates all of the undesired library members (**Fig. 1B**).^14^ Exemplified by the ease with which antibiotic resistance genes can be evolved in the laboratory,^15^ selections remove much of the time, cost and technical challenges associated with assaying vast libraries. However, transformations of interest for biotechnological applications rarely provide a growth advantage to a producing organism.

Successful *in vivo* selections thus rely on linking the host organism’s fitness to the genotype of an enzyme of interest, with the cell acting as both segregated compartment and expression host.^13,14^ Such a genotype-phenotype link can be achieved either via *genetic* or *chemical complementation* (**Figs. 1C**). In genetic complementation, the inability of the host organism to synthesize an essential metabolite (auxotrophy) is exploited to identify protein catalysts that are able to yield this compound. As a result, these efforts are restricted to a small set of enzymes for which an auxotrophic host is available or can be constructed by deleterious mutagenesis.^16-21^ In chemical complementation, an enzyme’s activity is linked to the production of a genetically-encoded reporter that enables cellular survival (**Fig. 1C**).^22^ By often relying on antibiotic resistance markers as reporter genes, this type of complementation avoids the use of auxotrophic strains and can theoretically be applied to a wider repertoire of enzymatic reactions. Traditionally, such selections function by small-molecule products of an enzymatic transformation binding either to a transcription factor or a genetically-encoded riboswitch, thereby regulating the transcription or translation of the employed reporter gene.^23,24^ More recently, linking enzymatic activities to phage propagation has resulted in a means to evolve enzymes autonomously along extended evolutionary trajectories.^25,26^

Additionally, a form of chemical complementation is employed when engineering orthogonal translation systems (OTSs) with the ability to expand the genetic code of organisms (**Figs. 1D**).^27^ Specifically, the activity of aminoacyl tRNA synthetase (aaRS) variants to load suppressor tRNAs with non-canonical amino acids (ncAAs) is routinely assessed by the suppression of an in-frame stop codon within an antibiotic resistance gene, such as a β-lactamase (**Fig. 1E**).^28,29^ Improved aaRS/tRNA pairs are therefore readily identified as they allow *Escherichia coli* to grow under higher concentrations of a β-lactam antibiotic (**Fig. 1E**). Notably, we recently demonstrated that this link between the activity of an OTS and host survival in presence of ampicillin can serve as a readout of unrelated catalytic activities.^30^ Specifically, we constructed a biocontainment strategy, in which new-to-nature reactions gated bacterial growth by transition-metal complexes that promote the formation of ncAAs.

In analogy, we surmised that the ncAA-dependency of recoded organisms could also serve as a starting point for engineering enzymes that can yield these essential building blocks from appropriate precursors. Here we provide proof-of-concept for such a chemical complementation strategy by demonstrating the *in vivo* directed evolution of carbamoylases, an enzyme class relevant for the biotechnological production of enantiopure (un)natural amino acids^.31^ In our directed evolution platform, the growth rates of *E. coli* correlated with the activity of carbamoylase variants they produce. Our strategy enabled us to identify enzymes that display turnover frequencies and catalytic proficiencies more than five orders of magnitude higher than those observed for the wild-type carbamoylase. Lastly, we also demonstrate that differences in growth rates provide a means to select improved variants from populations by performing serial passages under selective conditions. Given the ease with which improved carbamoylases were identified within our platform, we surmise that the use of recoded organisms in chemical complementation systems should facilitate the in vivo and continuous directed evolution of diverse biocatalysts.

## Results and Discussion

### Design and assembly of the chemical complementation system

Our directed evolution platform is based on the complementation of ncAA-dependent *E. coli* cells with enzymes that yield these essential compounds from appropriate precursor molecules (**Fig. 2A**). Establishing this functional link between enzymatic activity and survival requires the introduction of three genetic components: (1) an enzyme able to convert an appropriate precursor to the ncAA (=input); (2) an OTS selective for this ncAA (=sensor); and (3) a β-lactamase that can degrade carbenicillin (=readout). To avoid false positives that can result from the spontaneous mutation of the in-frame stop codon within the resistance gene, we make use of the engineered β-lactamase variant, TEM-1.B9,^32^ whose activity to degrade carbenicillin is strictly dependent on the incorporation of certain ncAAs, such as *L*-3-iodo- or *L*-3-nitro-tyrosine (3iY or 3nY). Following the production of all three components in *E. coli*, varying the concentration of carbenicillin should provide a tunable *selection pressure*, as more active enzymes should be able to proliferate in presence of higher antibiotic concentrations.

**Figure 2:**
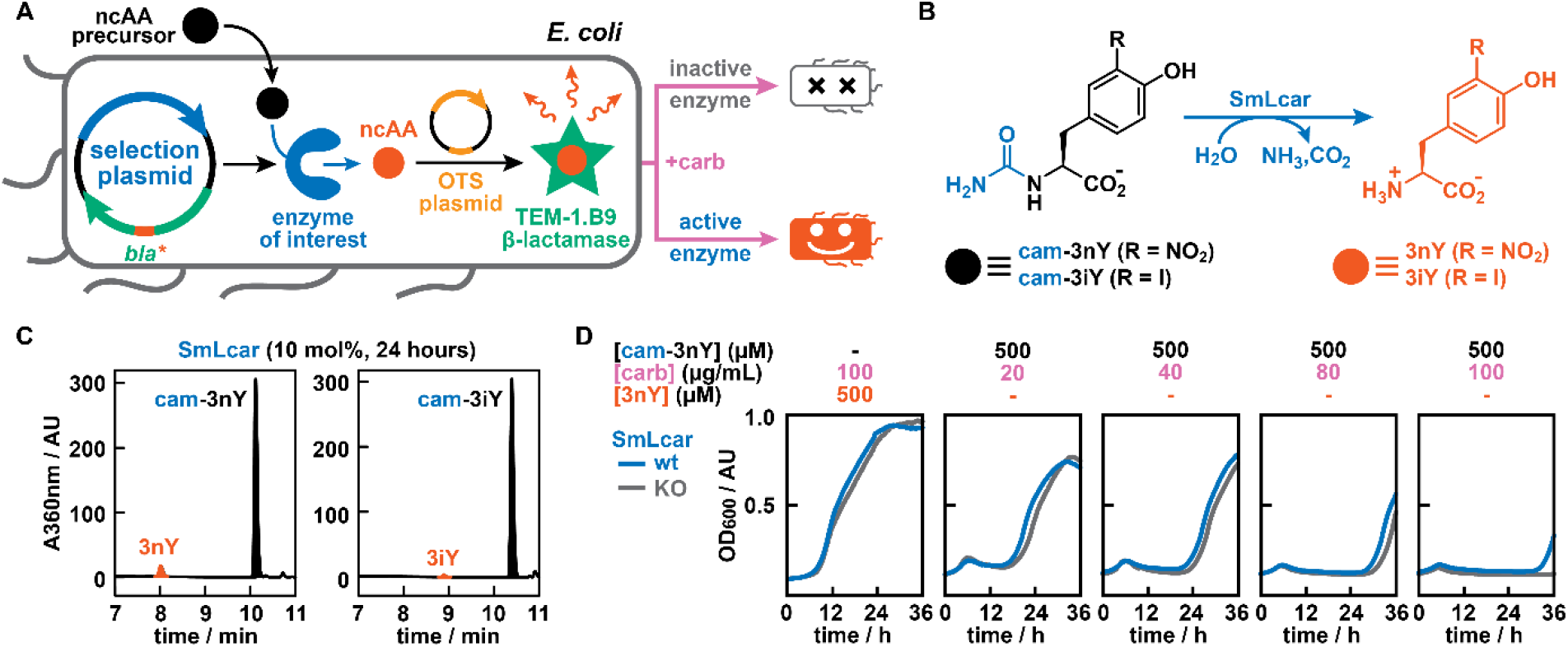
Design and validation of an in vivo enzyme engineering platform. **A:** Blueprint of a chemical complementation system based on recoded bacteria. A precursor is converted by an enzyme of interest to yield a ncAA, which is incorporated into a β-lactamase. Active enzymes should allow *E. coli* to grow in presence of carbenicillin, while inactive ones perish under the selection pressure **B:** Model transformations catalyzed by SmLcar that give rise to the ncAAs 3-nitro-_*L*_-tyrosine (3nY) and 3-iodo-_*L*_-tyrosine (3iY). **C:** HPLC chromatograms showing SmLcar is a weakly active catalyst for the conversion of carbamoylated ncAAs. **D:** Growth curves of *E. coli* in presence of varying concentrations of carbenicillin and 3nY or cam-3nY. The weak activity of SmLcar for the ncAA precursor provides a small growth advantage in the complementation system, when compared to an inactive enzyme variant (SmLcar_KO).

As a model biocatalyst class, we selected carbamoylases, which promote the hydrolytic cleavage of *N*-carbamoyl-amino acids (**Fig. 2B**).^31^ Together with hydantoinases, they make up a scalable and widely-employed enzymatic cascade for the production of enantiopure (un)natural amino acids (=hydantoin process).^33,34^ Based on its reportedly moderate activity for *N*-carbamoylated-*L*-tyrosine, we produced and purified SmLcar, a carbamoylase from *Sinorhizobium meliloti* strain CECT 4114.^35,36^ When challenged with the synthetic ncAA precursor *N*-carbamoyl-_*L*_-3nY (cam-3nY, 2 mM), SmLcar (10 μM) yielded 6.1% of the ncAA after 24 hours at pH 6 (**Fig. 2C**). Conversely, *N*-carbamoyl-_*L*_-3iY (cam-3iY) proved a more challenging substrate, with SmLcar failing to yield detectable levels of 3iY at pH 6 and only providing 0.8% of the ncAA at pH 8 under otherwise identical conditions.

A weakly active biocatalyst in hand, we assembled our chemical complementation system by introducing a selection and an OTS plasmid in *E. coli* (**Fig. 2A**). Both TEM-1.B9 and SmLcar were placed on a modified pACYCDuet-1 backbone (pACYC_GG), which features two type-IIS restriction sites for the modular exchange of the target enzyme and/or the β-lactamase (see **Supplementary Information** for details). The OTS – here consisting of the 3iY-aaRS and a suppressor tRNA – is encoded on the established pULTRA plasmid system, which features a resistance marker and an origin of replication that are compatible with the second vector.^37^ Lastly, the expression of all three genetic components is dependent on the addition of isopropyl-β-D-1-thiogalactopyranoside (IPTG), thus enabling the timed transcription and/or translation of all elements in our chemical complementation system.

### Validation of the complementation strategy

Before attempting to boost the activity of SmLcar for the synthetic ncAA precursor cam-3nY, we aimed to verify whether bacterial survival in presence of carbenicillin was dependent on (1) the supply of 3nY or (2) the addition of cam-3nY in combination with the production of SmLcar. In these experiments, we monitored bacterial growth at 30 °C by following the optical density at 600 nm (OD_600_) over a period of 36 hours (see **Supplementary Information** for details). Performing these experiments in 96-well plates enabled us to record the growth curves of cultures under varying conditions in parallel. To pinpoint the differences between an active and inactive biocatalyst, we also prepared the knock-out variant SmLcar_KO, in which arginine 292, an essential residue for substrate recognition, is replaced with an alanine.^36^

As expected, *E. coli* cultures featuring the OTS and TEM-1.B9 displayed robust growth upon addition of 3nY (500 µM) and carbenicillin (up to 500 µg/mL) to the media (**Extended Data Fig. 1**). The observed lag times were independent of the antibiotic concentration and comparable to those observed for cultures lacking carbenicillin (∼5-8 hours). Conversely, the addition of cam-3nY (500 µM) to SmLcar or SmLcar_KO containing bacteria resulted in extended lag phases, whose lengths increased with higher carbenicillin concentrations (**Fig. 2D**). We ascribe the observed proliferation of *E. coli* harboring the inactive variant SmLcar_KO to a combination of (1) the hydrolysis of cam-3nY catalyzed by endogenous enzymes with weak promiscuous activities for this synthetic substrate and (2) the (spontaneous) hydrolysis of carbenicillin over prolonged incubation times. Notably though, cells producing SmLcar consistently displayed somewhat shorter lag times than those featuring the knock-out variant, a trend that became more pronounced at higher carbenicillin concentrations (**Fig. 2D**). While the growth advantage provided by the active carbamoylase is small, we anticipated that the production of SmLcar variants featuring beneficial mutations should result in *E. coli* strains displaying significantly shorter lag times.

### Directed evolution of SmLcar

To identify positions in SmLcar that can be targeted by mutagenesis to boost the activity for cam-3nY, we predicted the structure of SmLcar using *AlphaFold2* (**Fig. 3A**, see **Supplementary Information** for details).^38^ Based on the available structural and functional information for carbamoylases,^31^ we first identified residues presumed to coordinate the pair of bivalent metal ions (e.g. Zn^2+^, Ni^2+^, or Mn^2+^) and those critical for substrate binding. From there, we selected four positions, which are either in proximity to the catalytic site (Gln91) or line the predicted binding pocket (Val150, Leu217, and Phe329, **Fig. 3A**). Libraries targeting each position were constructed using primers featuring degenerate NNK codons and 84 variants per residue were evaluated in parallel by following the OD_600_ in a plate reader in presence of 500 µM cam-3nY and 100 µg/mL carbenicillin (**Fig. 3B**, see **Supplementary Information** for details).

**Figure 3:**
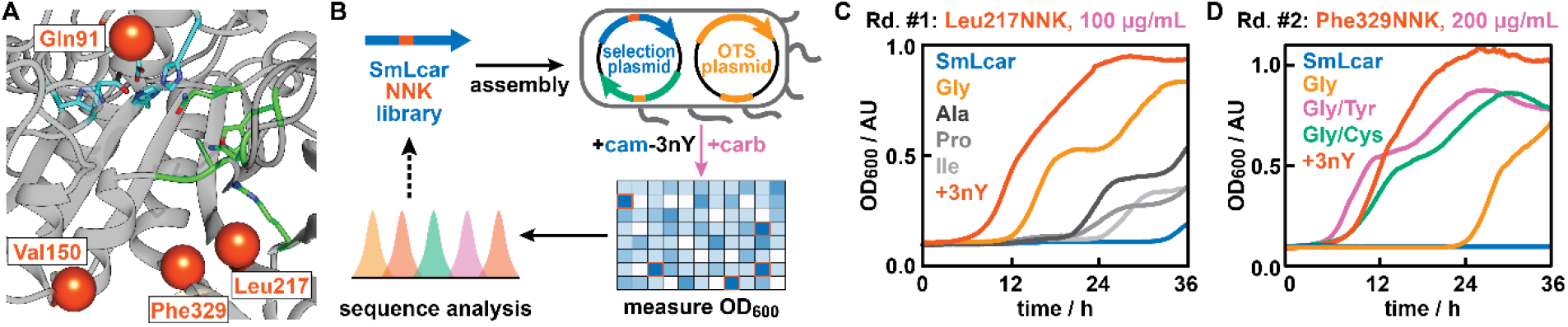
Directed evolution of SmLcar. **A:** Close-up view of the active site of SmLcar. Red spheres indicate positions which were targeted by site-directed mutagenesis in the evolution. Residues critical for binding of the two bivalent metal ions are highlighted in blue and amino acids that are either conserved or involved in substrate binding are shown in green. **B:** Workflow for the directed evolution of SmLcar in 96-well plates. **C:** Growth curves of enzyme variants with substitutions at Leu217 that provided a growth advantage in the first round of evolution. **D:** Growth curves of improved SmLcar variants featuring an additional mutation at Phe329. Note that increased carbenicillin concentrations result in longer lag times for SmLcar_G and the wild-type being unable to proliferate. Entries specified as +3nY refer to a positive control where cells producing SmLcar grow in presence of the ncAA (500 µM) instead of its synthetic precursor.

Gratifyingly, for the library targeting position Leu217, we detected several *complementation events*, as judged by the presence of significantly higher cell densities after 27 hours than those observed for the wild-type protein (**Extended Data Fig. 2**). Sequencing the SmLcar gene on their selection plasmids revealed that these variants featured common amino acid substitutions (Gly, Ala, Pro, or Ile), with the three variants displaying the highest OD_600_ values all being Leu217Gly. Inspecting the growth curves of individual variants over 36 hours further attests that all complementing clones displayed lag times that are shorter than those observed for the wild-type enzyme (**Fig. 3C** and **Extended Data Fig. 2**), with the glycine variant, SmLcar_G, again proving to be the fittest (lag time of ∼12 hours). Curiously, we observed a biphasic growth curve upon complementation for all variants. We cautiously ascribe this phenomenon to cells entering a second stationary phase after an initial growth-phase as cell division dilutes the produced 3nY and/or β-lactamase. At this stage, high concentrations of carbenicillin again prevent cell-wall synthesis, resulting in a temporary, second lag-phase.

Next, we selected SmLcar_G as a template for an additional round of site-directed mutagenesis, in which we targeted the remaining three positions for randomization (**Fig. 2A**). While the library of Gln91 did not result in a fitness increase, randomization of both Val150 and Phe329 yielded a number of hits that displayed faster growth than SmLcar_G (**Extended Data Fig. 3**). We selected the best-performing clones for each position and subjected them in triplicates to a rescreen under higher carbenicillin concentration (200 µg/mL, **Extended Data Fig. 4**). Consistent with a more stringent selection pressure, lag times for SmLcar_G increased to ∼24 hours, while the wild-type enzyme was unable proliferate under these conditions. Conversely, the rescreen identified two variants featuring either a Phe329Tyr or Phe329Cys substitution with lag times of only ∼8 hours and a somewhat less-pronounced second stationary phase (**Fig. 3D** and **Extended Data Fig. 4**). In fact, *E. coli* producing the engineered carbamoylases SmLcar_GY or SmLcar_GC displayed growth rates and lag times that are comparable to those observed upon direct supplementation of 3nY to recoded bacteria.

### Characterization of improved SmLcar variants

To demonstrate that shorter lag times of *E. coli* correlate with higher activities of SmLcar variants identified in the screen, we produced and purified these carbamoylases and evaluated their performance on ncAA precursors in vitro. While the wild-type enzyme at 10 μM (0.5 mol%) yielded a modest 6.1% of the ncAA from cam-3nY (500 μM) over 24 hours (**Fig. 2A**), all variants identified in the evolution campaign were able to fully convert the synthetic precursor under these conditions. Shortening the reaction time to 1 hour and lowering the enzyme concentration 20-fold to 0.5 μM revealed that lag-times in 96-well plates accurately report on the activity of engineered carbamoylase variants. Specifically, SmLcar_G, the best performing variant in the first round, and the double mutants SmLcar_GC and SmLcar_GY gave 3nY in 4.5%, 38.8% and 50.5% yield, respectively (**Fig. 4A**).

**Figure 4:**
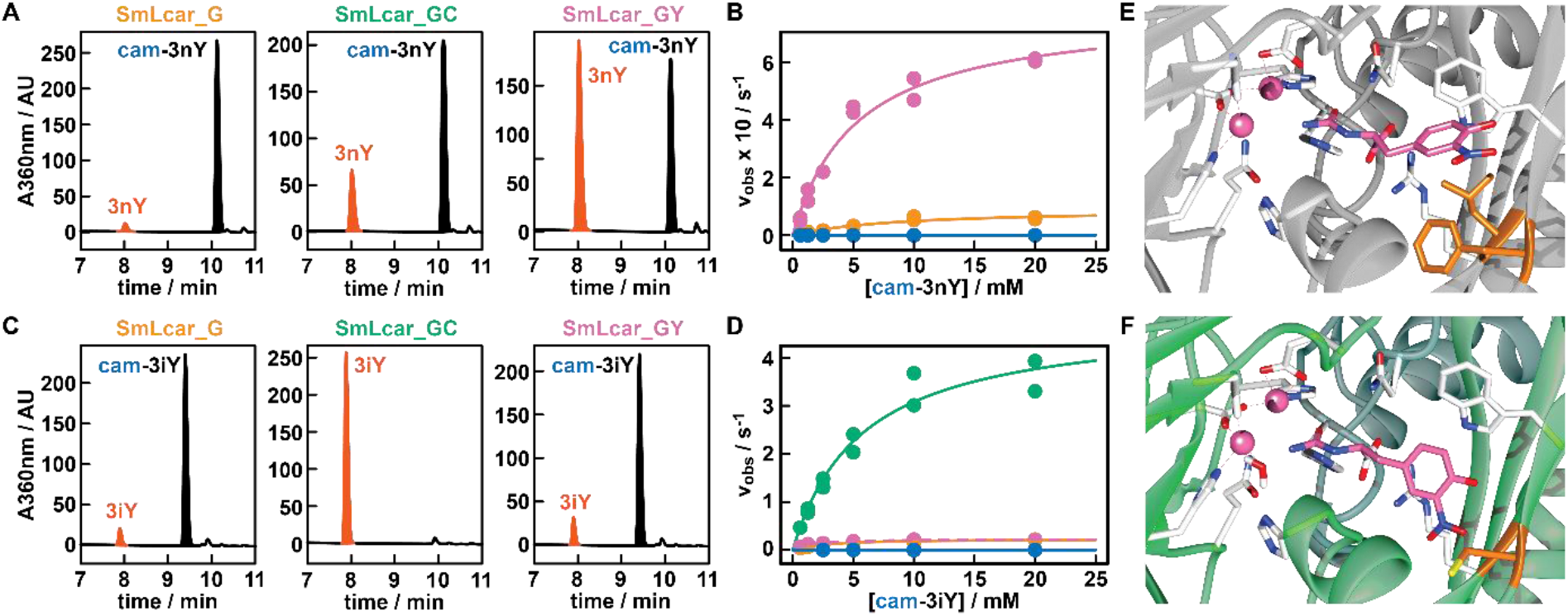
Characterization of improved SmLcar variants: **A:** HPLC chromatograms showing the activities of engineered SmLcar variants for cam-3nY. **B:** Saturation kinetics for SmLcar (blue), SmLcar_G (orange) and SmLcar_GY (purple) for cam-3nY. The corresponding kinetic parameters obtained from fitting data points to the Michaelis Menten equation are listed in **Table 1. C:** HPLC chromatograms showing the activities of engineered SmLcar variants for cam-3iY. **D:** Saturation kinetics for SmLcar (blue), SmLcar_G (orange), SmLcar_GY (purple), and SmLcar_GC (green) for cam-3iY. The corresponding kinetic parameters obtained from fitting data points to the Michaelis Menten equation are listed in **Table 1. E-F:** Close-up view of the active sites of SmLcar (**E**, PDB: 8APZ) and SmLcar_GC (**F**, PDB: 8AQ0) with cam-3nY docked into the obtained crystal structures.

Next, we determined the kinetic parameters of SmLcar and the best variants from each round to accurately pinpoint their level of improvement (**Fig. 4B** and **Table 1**). As expected, the parent enzyme displayed a low turnover frequency (*k*_cat_ = 1.05×10-4 ± 1.01×10-5 s-1) and catalytic proficiency (*k*_cat_/*K*_M_ = 3.54 ± 0.98 × 10-2 M-1 s-1). Notably, the single Leu219Gly substitution in SmLcar_G provided a >850-fold boost in turnover frequencies (*k*_cat_ = 0.093 ± 0.014 s-1) and resulted in an ∼300-fold increase in catalytic proficiencies (*k*_cat_/*K*_M_ = 10.4 ± 2.6 M-1 s-1). The addition of the Phe329Tyr substitution in SmLcar_GY also proved beneficial, providing a further 8.4- and 14.2-fold increase in *k*_cat_ (=0.78 ± 0.05 s-1) and *k*_cat_/*K*_M_ (=148 ± 12.9 M-1 s-1), respectively.

**Table 1.** Kinetic parameters of SmLcar and engineered variants for the ncAA precursors cam3nY and cam3iY

| Enzyme | cam-3-L-nitrotyrosine (cam-3nY) |  |  |  | cam-3-L-iodotyrosine (cam-3iY) |  |  |  |
| --- | --- | --- | --- | --- | --- | --- | --- | --- |
| | $k_{\text{cat}} (\text{s}^{-1})$ | $K_{\text{M}} (\text{mM})$ | $k_{\text{cat}}/K_{\text{M}} (\text{M}^{-1} \text{s}^{-1})$ | relative | $k_{\text{cat}} (\text{s}^{-1})$ | $K_{\text{M}} (\text{mM})$ | $k_{\text{cat}}/K_{\text{M}} (\text{M}^{-1} \text{s}^{-1})$ | relative |
| SmLcar | $1.05 \times 10^{-4}$<br>($1.01 \times 10^{-5}$) | 2.98<br>(0.87) | $3.54 \times 10^{-2}$<br>($9.77 \times 10^{-3}$) | 1 | $6.92 \times 10^{-5}$<br>($2.09 \times 10^{-5}$) | 13.8<br>(7.91) | $5.00 \times 10^{-3}$<br>($3.15 \times 10^{-3}$) | 1 |
| SmLcar_G | $9.29 \times 10^{-2}$<br>($1.36 \times 10^{-2}$) | 8.91<br>(2.81) | 10.4<br>(2.58) | 294 | 0.232<br>(0.013) | 2.58<br>(0.46) | 89.9<br>(13.2) | $1.80 \times 10^4$ |
| SmLcar_GY | 0.78<br>(0.05) | 5.30<br>(0.88) | 148<br>(12.9) | 4,170 | 4.84<br>(0.39) | 5.71<br>(1.14) | 849<br>(127) | $1.70 \times 10^5$ |
| SmLcar_GC | n.d. | n.d. | n.d. | n.d. | 0.231<br>(0.05) | 1.45<br>(0.16) | 159<br>(14.2) | $3.18 \times 10^4$ |

Curious whether the substitutions identified in the directed evolution present a more general solution for the conversion of *m*-substituted tyrosine analogs, we determined the activity of engineered carbamoylases for cam-3iY. Indeed, SmLcar_G (10 μM) was able to fully convert the synthetic precursor (500 μM) within 24 hours at pH 8, while the wild-type enzyme gave only 0.8% yield under identical conditions. To obtain a representative comparison between our evolved carbamoylases, we again lowered the enzyme concentration to 0.5 μM and shortened reaction times to 1 hour (**Fig. 4C**). Surprisingly, SmLcar_G and SmLcar_GY gave rise to comparable yields (7.3% and 8.0%, respectively), indicating that the improved activity of the tyrosine variant for cam-3nY reflects a certain specialization toward the screening substrate. Unexpectedly, the Phe329Cys substitution resulted in significantly improved reactivity, as SmLcar_GC fully converted cam-3iY under these conditions (**Fig. 4C**).

Determining the steady-state kinetic parameters confirmed the increased activity of all variants for cam-3iY, when compared to the wild-type (*k*_cat_ = 6.92×10-5 ± 2.09×10-5 s-1; *k*_cat_/*K*_M_ = 5.00×10-3 ± 3.15×10-3 M-1 s-1). The Leu217Gly substitution again proved crucial and boosted the turnover frequency (=0.23 ± 0.01 s-1) and catalytic efficiency (89.9 ± 13.2 M-1 s-1) by more than three and four orders of magnitude, respectively (**Fig. 4D** and **Table 1**). Consistent with the conversions obtained by HPLC, the addition of Phe329Tyr resulted only in a minor improvement in catalytic proficiency (=159 ± 14.2 M-1 s-1), when compared to SmLcar_G. In stark contrast, installing a cysteine at position 329 proved highly beneficial, boosting the turnover number (4.84 ± 0.39 s-1) and the catalytic efficiency (849 ± 127 M-1 s-1) by another order of magnitude. When compared to the parent carbamoylase, SmLcar_GC displays a ∼70,000-times higher *k*_cat_ and a ∼170,000-improvement of *k*_cat_/*K*_M_. Combined, these results attest that our strategy of linking unrelated enzymatic activities to bacterial proliferation presents an effective means to engineer weakly active biocatalysts that do not fulfill a metabolic role.

### Crystal structures of SmLcar and SmLcar_GC

To rationalize the massive performance improvements of our engineered carbamoylases, we obtained crystal structures for SmLcar and the double mutant SmLcar_GC (**Extended Data Fig. 5** and see **Supplementary Information** for details). Guided by the electron densities of unknown small molecules bound to the active sites of SmLcar and SmLcar_GC (**Extended Data Fig. 6**), we performed docking studies (*YASARA*)^39^ with the synthetic precursor, cam-3nY, for both enzymes. Comparing the resulting structures of the wild-type and the evolved variant revealed a striking difference in the orientation of the substrate in their respective binding pockets (**Figs. 4E-F**). Specifically, the critical Leu217Gly substitution identified in the first round enlarges the binding pocket and allows the aromatic side chain to flip by ∼90°. While our kinetic characterization reveals that this does not drastically alter the affinity of the ncAA precursor for the enzyme, as judged by the small differences in *K*_M_, this conformation appears to be highly beneficial for the attack of water at the adjacent metal-binding site. Lastly, position 329 is distal to Gly217 and the introduced cysteine in SmLcar_GC (or tyrosine in SmLcar_GY) interacts with the substrate to likely further lock it in an active conformation.

### Recapitulating the evolution of SmLcar_GY on the population level

In developing a robust, chemical complementation strategy we aimed to facilitate the *selection* of improved biocatalysts from diverse populations. Overcoming the need of assessing the activity of enzyme variants one-by-one is highly desirable, as selections combine a high throughput with operational simplicity.^13,14^ In our platform, the fitness of *E. coli* in presence of carbenicillin and cam-3nY is dictated by the activity of the produced carbamoylase variant. With improved biocatalysts resulting in shorter lag times under these selective conditions, *E. coli* harboring such variants should outcompete bacteria featuring less active carbamoylases. Relying on this robust genotype-phenotype link, we set out to provide proof-of-concept that our platform is amenable to elicit improved enzymes from populations based on the growth advantage they bestow on a producing organism.

Toward this end, we first performed a mock selection by mixing cultures containing the wild-type carbamoylase and SmLcar_GY in a 1:1 ratio (**Fig. 5A** and **Extended Data Table 1**). Isolating the selection plasmid and analyzing the carbamoylase gene encoded on it by Sanger sequencing allows us to (1) unambiguously assign either variant, (2) estimate their relative frequency within a given a population, and (3) track changes in the population over time. Prior to selection, the mixed population (OD_600_ ≈ 0.01) reflected the anticipated 1:1 ratio between the wild-type and SmLcar_GY, which was subsequently challenged to grow overnight in presence of cam-3nY (500 μM) and carbenicillin (50 μg/mL). Sequencing the plasmids isolated from the densely-grown culture the next day revealed that the population had become dominated by SmLcar_GY (**Fig. 5A**). As such, this result attests that the growth advantage provided by a more active carbamoylase under selection conditions enables its amplification in a mixed population.

**Figure 5:**
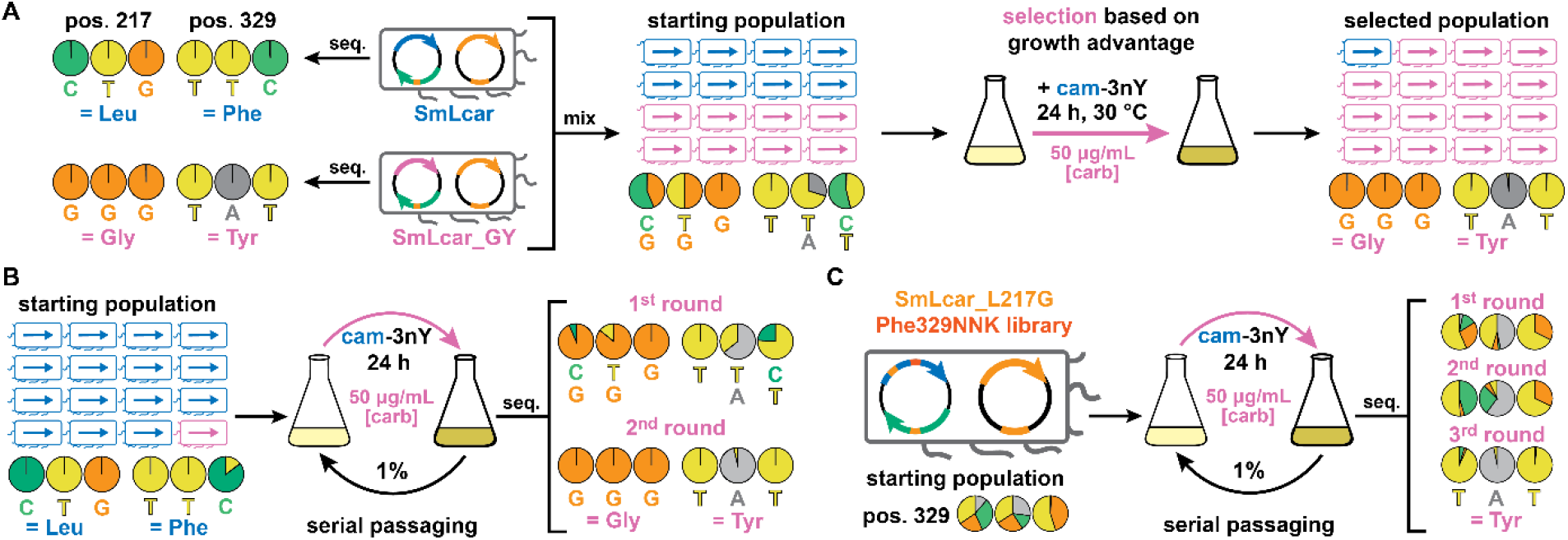
Selection of improved carbamoylases on the population level. **A:** Mock selection containing an equal mixture of SmLcar and SmLcar_GY. Overnight growth under selective conditions results in the improved carbamoylase becoming the dominant species. Base calls from Sanger sequencing of individual variants or populations are displayed as pie charts. **B:** Mock selections for a ≥10:1 mixture of SmLcar and SmLcar_GY. Cultures grown overnight are subjected to serial passaging under selective pressure. Two such growth-dilution cycles are sufficient to yield a population that is dominated by SmLcar_GY **C:** Application of the selection scheme by serial passaging to the second-round SmLcar_G_Phe329NNK library. Starting from a diverse library, three growth-dilution cycles elicit SmLcar_GY as the dominant variant as judged by Sanger sequencing.

To test the ability of our platform to identify better biocatalysts that are less abundant, we performed another mock selection, in which wild-type and SmLcar_GY cultures were mixed in a ≥10:1 ratio (**Fig. 5B**). Sequencing the starting population failed to detect appreciable levels of the improved carbamoylase, a fact that likely reflects the limitations of Sanger sequencing. This heavily-skewed population was subjected to *serial passaging*, that is cultures were diluted by a factor of 100 following growth under selection conditions for 24 hours. Again, we traced the fate of the population by sequencing plasmids recovered after each growth period. Strikingly, within 24 hours SmLcar_GY became the dominant variant in the population, with an additional dilution-growth step resulting in a population devoid of the wild-type carbamoylase (**Fig. 5B** and **Extended Data Table 1**). To exclude the possibility of SmLcar_GY being amplified independent of its catalytic activity, we also performed serial dilutions on the mixed population by adding 3nY to the growth media. As expected, in absence of selecting for catalytic activity, sequencing cultures following two dilution-growth cycles did not show a significant change in the population (**Extended Data Fig. 7**).

Combined, the amplification factors observed for SmLcar_GY in the mock selections attest on a robust genotype-phenotype link that allows individual cells rather than the whole population to benefit from producing a more proficient enzyme. Consequently, *E. coli* cells harboring less active carbamoylases cannot strongly exploit escape mechanisms, such as beginning to divide once carbenicillin has been (largely) degraded in the media or taking up 3nY that is released by fitter bacteria. Avoiding these unintended amplification and escape mechanisms is critical for the successful adaptation of our platform to select improved carbamoylases from libraries.

Encouraged by these observations, we challenged our platform to elicit SmLcar_GY from the SmLcar_G_Phe329NNK library we used in the second round of evolution in 96-well plates (**Fig. 5C**). Unlike in mock selections, where we employ the far inferior wild-type carbamoylases, the activity differences in this population are significantly smaller, with SmLcar_G and SmLcar_GC being about 10% and 50% as active as the desired tyrosine variant (*cf*. **Fig. 4A**). Sequencing the starting population attested on an equal distribution of all desired nucleotides for the NNK stretch that encodes for all 20 amino acids at position 329. As before, we performed serial passages under selective and non-selective conditions and followed changes in the population by sequencing. As expected, the culture grown in presence of 3Ny did not undergo significant changes following two growth-dilution cycles (**Extended Data Fig. 8**). Conversely, the population subjected to selective conditions was dynamic, with the codon TAT corresponding to tyrosine ultimately dominating the population following three passages (**Fig. 5C** and **Extended Data Table 1**). To independently verify this result, we sequenced ten individual colonies from the population and found that six of them indeed encoded for SmLcar_GY (**Extended Data Table 2**). Lastly, applying the same selection mechanism to the first-round library SmLcar_Leu217NNK also elicited SmLcar_G as the dominant variant following three growth-dilution cycles (**Extended Data Fig. 8**).

## Conclusions

Utilizing in vivo selections for the directed evolution of biocatalysts promises to quantitatively capture enzyme activity and efficiently isolate desired variants from diverse populations.13,14 Adopting powerful in vivo selections for enzymes that do not fulfill a metabolic function requires the introduction of genetically-encodable elements that link cellular survival to a transformation of interest.24 Here, we demonstrate that ncAA-dependent organisms represent an ideal platform for constructing such chemical complementation systems that can be employed for enzyme engineering purposes. Specifically, we showcase that bacterial proliferation is dependent on an enzyme that can provide the ncAA from an appropriate precursor and that the growth advantage provided by variants is dependent on their activities.

Using this artificial link between enzyme activity and bacterial proliferation, we evolved the carbamoylase SmLcar to accept carbamoylated *m*-substituted tyrosine analogs. Using bacterial growth in presence of carbenicillin as the *only* readout allowed us to identify SmLcar variants that hydrolyzed ncAA precursors with catalytic proficiencies up to five orders of magnitude higher than those observed for the parent enzyme. While the initial evolution was performed in 96-well plates, the fact that our platform does not require any handling steps following inoculation significantly simplifies and streamlines the discovery of carbamoylases with improved kinetic parameters. Furthermore, we demonstrated that the growth advantage provided by improved carbamoylases is largely confined to producing cells and does not increase the fitness of less active variants in a population. This robust genotype-phenotype link enabled us to elicit improved carbamoylases in mock selections and recapitulate the evolution of SmLcar on the population level. These selections were performed by straightforward serial passaging following overnight growth, thus further minimizing the handling steps for the experimentalist. Moreover, the fact that our platform is not dependent on specialized and/or expensive equipment argues well for its widespread application.

In the future, we will explore the full capabilities of our chemical complementation system, which promises the following developments. First, building on well-established strategies to expand the genetic code of *E. coli*, our platform is applicable to any reaction that can be linked to the formation of one of the >150 non-canonical building blocks that have been incorporated into proteins in vivo.^28,29^ Critically, by having the potential to provide the same readout (=survival) for reactions catalyzed by mechanistically-diverse enzymes, our platform is not only innovative but also flexible in nature. Second, taking advantage of the operational simplicity and realizing the high throughput of biological selections will allow for a thorough interrogation of the available sequence space. As such, selecting better enzymes from large and diverse libraries should facilitate the identification of beneficial mutations that cannot be rationalized or exceedingly rare mutations that work in synergy.^6-8^ Lastly, the combination of our approach with means to introduce mutations in genes of interest in vivo,^26,40,41^ should allow for establishing continuous evolution approaches. By tailoring diverse enzymatic activities with minimal or without human intervention, such developments will allow us to explore long evolutionary trajectories in parallel.

## EXTENDED DATA

**Extended Data Figure 1:**
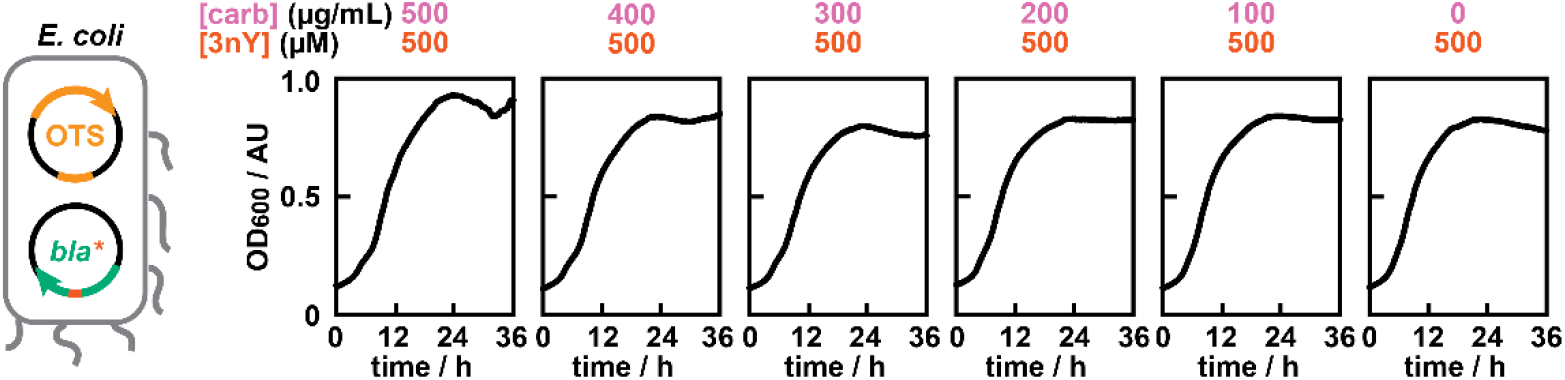
Growth curves of *E. coli* featuring the OTS and the β-lactamase, TEM-1.B9, in presence of varying concentrations of carbenicillin and fixed concentrations of 3nY. The observed lag times were independent of the selection pressure and comparable to those observed for cultures lacking carbenicillin.

**Extended Data Figure 2:**
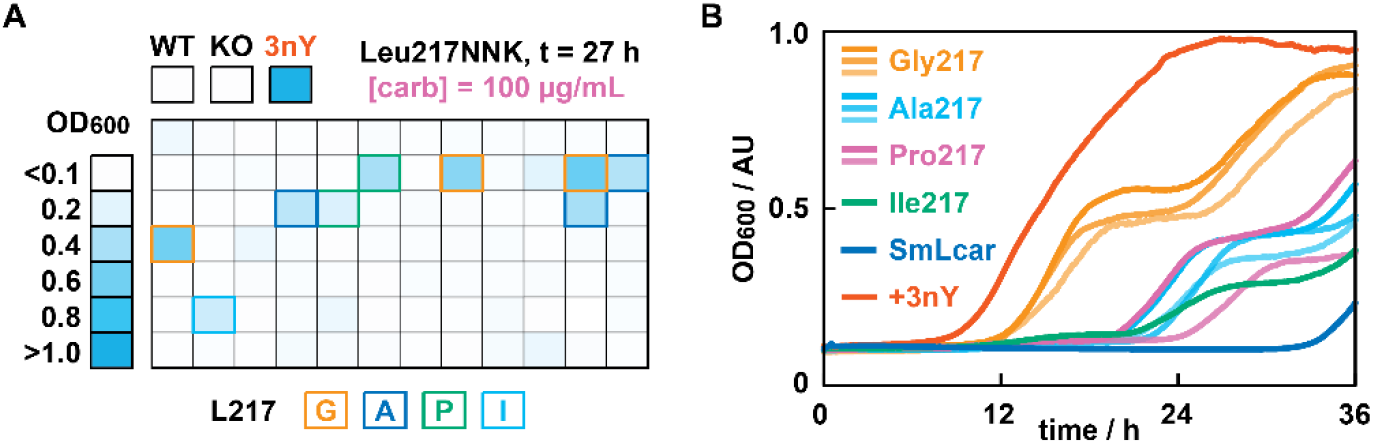
**A:** Observed cell densities (OD_600_) for SmLcar_Leu217NNK library after 27 hours of incubation in presence of 100 μg/mL carbenicillin. Variants that displayed significantly increased proliferation with respect to the parent enzyme were identified by sequencing. **B:** Extracted growth curves for hits identified in the 96-well plate screens. The Leu217Gly substitution consistently showed the shorted lag times.

**Extended Data Figure 3:**
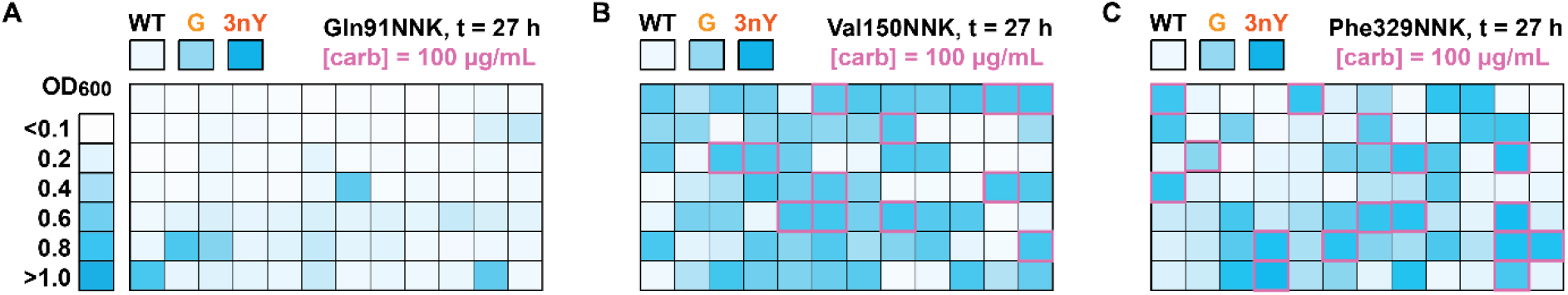
**A-C:** Observed cell densities (OD_600_) for SmLcar_G_NNK libraries targeting Gln91 (**A**), Val150 (**B**), and Phe329 (**C**). Variants displaying significantly improved proliferation with respect to SmLcar_G were selected for a rescreen (highlighted in pink) following manual inspection of individual growth curves.

**Extended Data Figure 4:**
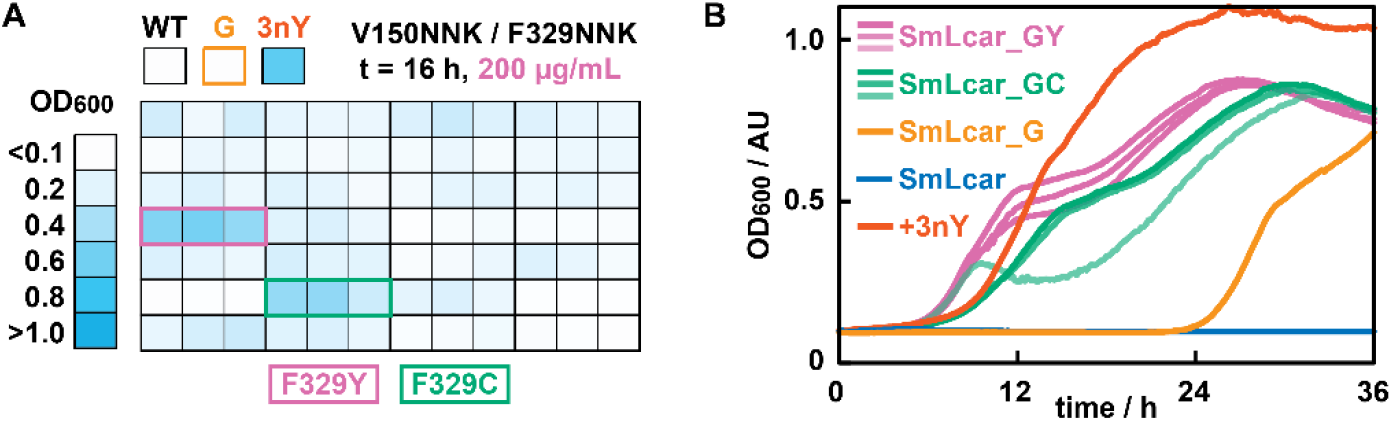
**A:** Observed cell densities (OD_600_) in the rescreen of SmLcar_G_NNK libraries after 16 hours of incubation in presence of 200 μg/mL carbenicillin. Variants were measured in triplicates and those performing best were identified by sequencing. **B:** Extracted growth curves for SmLcar_GY and SmLcar_GC, showing that the former consistently displayed the shortest lag times.

**Extended Data Figure 5:**
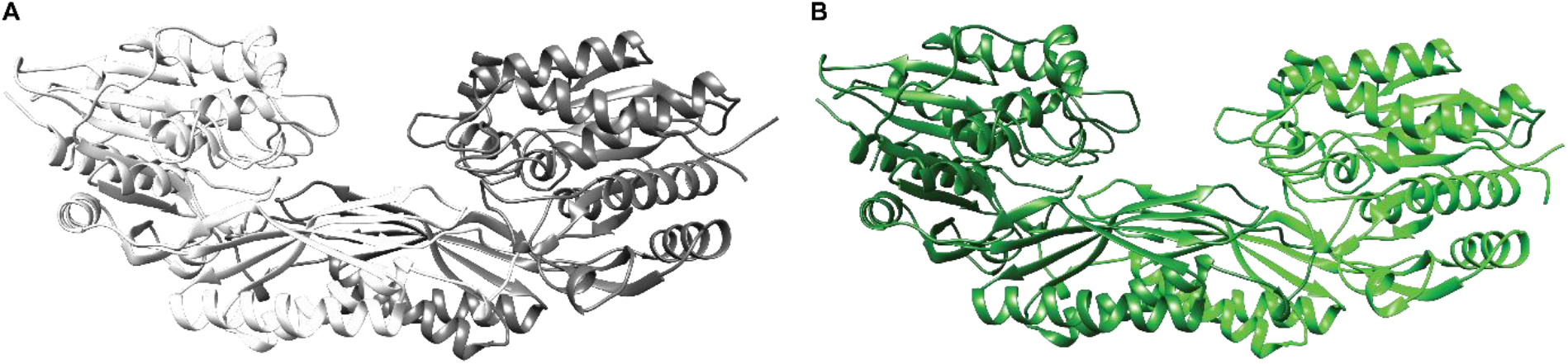
**A-B:** Crystal structures of SmLcar at 1.75 Å (**A**, PDB: 8APZ) and SmLcar_GC at 2.30 Å (**B**, PDB: 8AQ0).

**Extended Data Figure 6:**
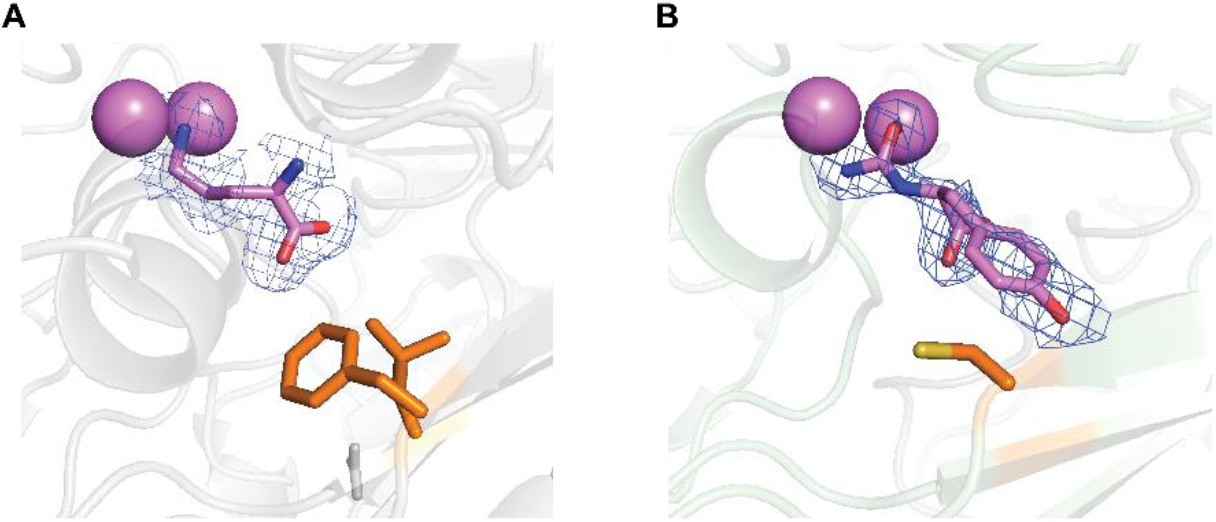
**A-B**. Close-up view of the active sites of SmLcar and SmLcar_GC. Polder omit maps contoured at 3σ for each of the complexes show the electron density of unknown molecules bound to the respective active sites. Based on the occupancy, we tentatively assigned these densities to ornithine (SmLcar) and carbamoyl-L-tyrosine (SmLcar_GC). Their structures and the two metal ions are shown in purple, while side chains of residues that changed in the evolution campaign are shown in orange.

**Extended Data Figure 7:**
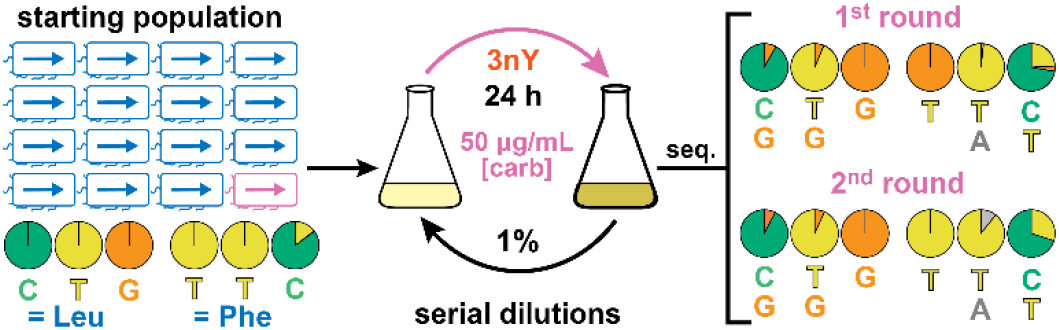
Subjecting a ≥10:1 mixture of SmLcar and SmLcar_GY in absence of a selection pressure does not result in the amplification of the improved carbamoylase. Note that the addition of 3nY (500 μM) to the growth media does not require carbamoylase activity for the production of the β-lactamase.

**Extended Data Figure 8:**
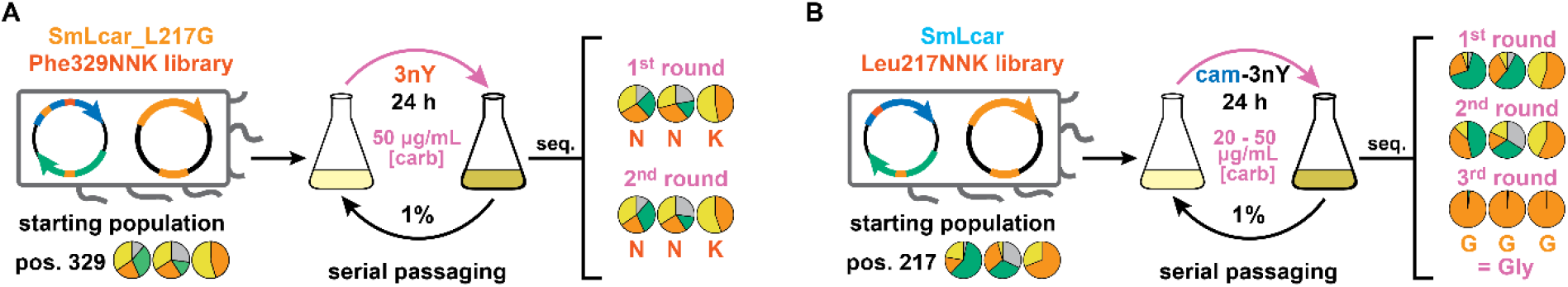
**A:** Performing serial passages with the SmLcar_G_Phe329NNK library in absence of a selection pressure does not result in significant changes in the population as judged by Sanger sequencing. **B:** Application of the selection scheme by serial passaging to the first-round SmLcar_Leu217NNK library. Starting from a diverse library, three growth-dilution cycles elicit SmLcar_G as the dominant variant as judged by Sanger sequencing. Note that the first growth-dilution cycle was performed at 20 μg/mL, a selection pressure that was likely too low to result in a significant change in the composition of the library.

**Extended Data Table 1.** Base calls extracted from Sanger sequencing traces for all selections performed in this study.

| SmLcar : SmLcar_GY = 1:1 |  |  |  |  |  |  |  |  |
| --- | --- | --- | --- | --- | --- | --- | --- | --- |
| Pos. | starting population |  |  |  | 1st overnight |  |  |  |
| 217 | A | C | G | T | A | C | G | T |
| 1 | 0.16 | 49.5 | 50.3 | 0 | 8.99 | 0 | 91 | 0 |
| 2 | 0.19 | 0 | 34 | 65.8 | 0.88 | 0 | 99.1 | 0 |
| 3 | 0 | 0 | 100 | 0 | 0 | 0 | 99.9 | 0.14 |
| 329 | A | C | G | T | A | C | G | T |
| 1 | 0.07 | 0.22 | 0 | 99.7 | 0.25 | 0.17 | 3.97 | 95.6 |
| 2 | 28.8 | 0.14 | 0 | 71 | 92.9 | 0 | 0 | 7.09 |
| 3 | 0.14 | 50.6 | 0.27 | 49 | 0.59 | 0 | 0.59 | 98.8 |

| SmLcar : SmLcar_GY = 10:1 no selection pressure |  |  |  |  |  |  |  |  |  |  |  |  |
| --- | --- | --- | --- | --- | --- | --- | --- | --- | --- | --- | --- | --- |
| Pos. | starting population |  |  |  | 1st overnight |  |  |  | 2nd overnight |  |  |  |
| 217 | A | C | G | T | A | C | G | T | A | C | G | T |
| 1 | 0 | 100 | 0 | 0 | 0 | 91.6 | 8.25 | 0.13 | 0 | 92 | 7.69 | 0.28 |
| 2 | 0 | 0 | 0.34 | 99.7 | 0 | 0.12 | 6.39 | 93.5 | 0 | 0 | 6.85 | 93.1 |
| 3 | 0 | 0 | 100 | 0 | 0 | 0 | 99.8 | 0.21 | 0 | 0 | 99.6 | 0.42 |
| 329 | A | C | G | T | A | C | G | T | A | C | G | T |
| 1 | 0 | 0.25 | 0 | 99.8 | 0 | 0.35 | 0 | 99.6 | 0 | 0.22 | 0 | 99.8 |
| 2 | 0 | 0.33 | 0 | 99.7 | 1.84 | 0.12 | 0 | 98 | 10.8 | 0.18 | 0 | 89 |
| 3 | 0 | 85.1 | 0.35 | 14.6 | 0 | 72.2 | 3.66 | 24.1 | 0 | 69.9 | 0.11 | 30 |

| SmLcar : SmLcar_GY = 10:1 |  |  |  |  |  |  |  |  |  |  |  |  |
| --- | --- | --- | --- | --- | --- | --- | --- | --- | --- | --- | --- | --- |
| Pos. | starting population |  |  |  | 1st overnight |  |  |  | 2nd overnight |  |  |  |
| 217 | A | C | G | T | A | C | G | T | A | C | G | T |
| 1 | 0 | 100 | 0 | 0 | 0 | 6.32 | 93.7 | 0 | 0 | 0.2 | 99.8 | 0 |
| 2 | 0 | 0 | 0.34 | 99.7 | 0 | 0 | 85.8 | 14.2 | 0.49 | 0 | 99.5 | 0 |
| 3 | 0 | 0 | 100 | 0 | 0 | 0 | 99.6 | 0.38 | 0 | 0 | 99.7 | 0.3 |
| 329 | A | C | G | T | A | C | G | T | A | C | G | T |
| 1 | 0 | 0.25 | 0 | 99.8 | 0.2 | 0.2 | 0 | 99.6 | 0.23 | 0.17 | 0 | 99.6 |
| 2 | 0 | 0.33 | 0 | 99.7 | 64.3 | 0.09 | 0 | 35.6 | 96.2 | 0 | 0 | 3.76 |
| 3 | 0 | 85.1 | 0.35 | 14.6 | 0.25 | 23.9 | 0.12 | 75.7 | 0.46 | 1.23 | 0.15 | 98.2 |

| SmLcar_Phe329_NNK library |  |  |  |  |  |  |  |  |  |  |  |  |  |  |  |  |
| --- | --- | --- | --- | --- | --- | --- | --- | --- | --- | --- | --- | --- | --- | --- | --- | --- |
| Pos. | starting population |  |  |  | 1st overnight |  |  |  | 2nd overnight |  |  |  | 3rd overnight |  |  |  |
| 329 | A | C | G | T | A | C | G | T | A | C | G | T | A | C | G | T |
| 1 | 11.3 | 31.8 | 22.2 | 34.6 | 3.13 | 12.8 | 27.5 | 56.5 | 0.48 | 44.9 | 3.97 | 50.6 | 0.27 | 4.43 | 2.3 | 93 |
| 2 | 27 | 13.8 | 25.1 | 34.1 | 46.8 | 2.02 | 5.84 | 45.3 | 60 | 27.4 | 5.39 | 7.23 | 96.4 | 0 | 0.41 | 3.18 |
| 3 | 0.09 | 0.81 | 44.3 | 54.8 | 0.1 | 0 | 32.3 | 67.6 | 0.16 | 0 | 32.1 | 67.8 | 0.23 | 0 | 1.71 | 98.1 |

| SmLcar_Phe329_NNK library no selection |  |  |  |  |  |  |  |  |  |  |  |  |  |
| --- | --- | --- | --- | --- | --- | --- | --- | --- | --- | --- | --- | --- | --- |
| Pos. | starting population |  |  |  | 1st overnight |  |  |  | 2nd overnight |  |  |  | 3rd overnight |
| 329 | A | C | G | T | A | C | G | T | A | C | G | T |  |
| 1 | 11.3 | 31.8 | 22.2 | 34.6 | 12.6 | 25.2 | 28.8 | 33.4 | 15.2 | 27 | 32.3 | 25.5 |  |
| 2 | 27 | 13.8 | 25.1 | 34.1 | 22.4 | 16.5 | 32.2 | 28.9 | 21.5 | 19.3 | 30.3 | 28.8 |  |
| 3 | 0.09 | 0.81 | 44.3 | 54.8 | 0.09 | 1.12 | 46.6 | 52.2 | 0.09 | 0 | 40.9 | 59 |  |

| SmLcar_Leu217_NNK library |  |  |  |  |  |  |  |  |  |  |  |  |  |  |  |  |
| --- | --- | --- | --- | --- | --- | --- | --- | --- | --- | --- | --- | --- | --- | --- | --- | --- |
| Pos. | starting population |  |  |  | 1st overnight |  |  |  | 2nd overnight |  |  |  | 3rd overnight |  |  |  |
| 217 | A | C | G | T | A | C | G | T | A | C | G | T | A | C | G | T |
| 1 | 3.22 | 58.8 | 15.5 | 22.5 | 4.24 | 65.7 | 24.7 | 5.4 | 0.28 | 46.4 | 40.3 | 13 | 0 | 1.72 | 98.3 | 0 |
| 2 | 32 | 31.7 | 31.7 | 4.53 | 7.82 | 53.7 | 30.6 | 7.82 | 33.9 | 31.7 | 16.4 | 18 | 1.25 | 0 | 98.8 | 0 |
| 3 | 0 | 0 | 69.1 | 30.9 | 0.58 | 4.19 | 52.7 | 42.5 | 0.09 | 0 | 57.2 | 42.7 | 0 | 0 | 100 | 0 |

**Extended Data Table 2.**
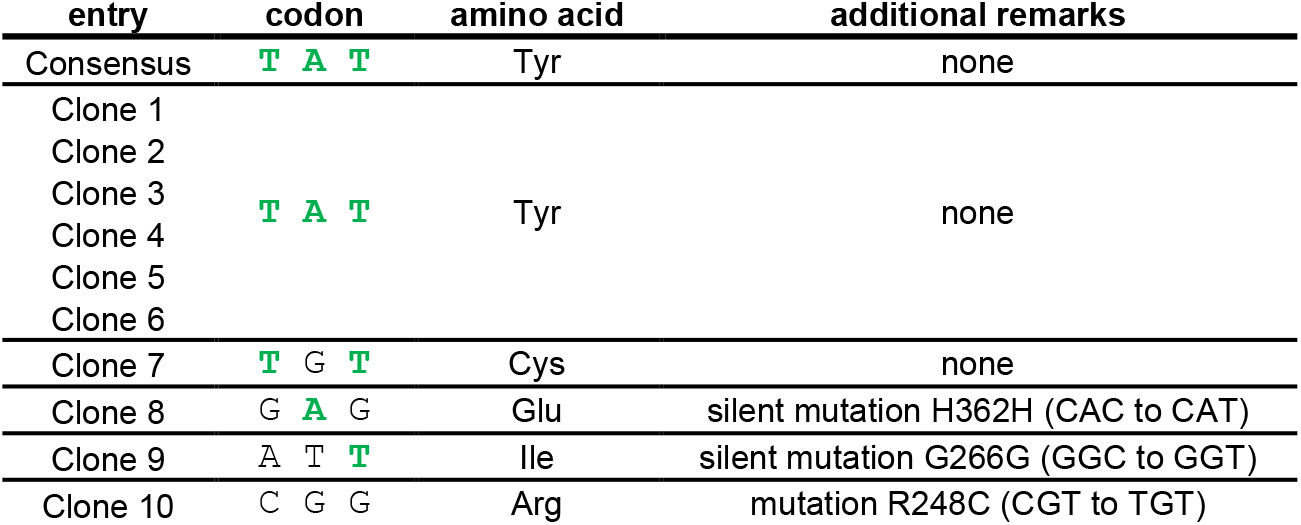
Sequencing results for 10 clones isolated after three growth-dilution cycles with the Phe329NNK library. Codons and corresponding amino acids are given for position 329, with additional remarks listed. Nucleotides matching the consensus sequence are highlighted in green.

## Acknowledgements

C.M. and R.R. are thankful to A. Boverio for the help with the protein crystallization, to A.M.W.H. Thunnissen for the help with AlphaFold2, and to D. Sauer for the guidance with the docking simulations. C.M. acknowledges the NOW (VENI grant 722.017.007 and ENW-M grant OCENW.M20.278) for funding.

## Notes

### Competing Interest Statement

The authors have declared no competing interest.

## References

1 Knowles, J. R. Enzyme catalysis: not different, just better. Nature 350, 121–124 (1991).

2 Wolfenden, R. & Snider, M. J. The Depth of Chemical Time and the Power of Enzymes as Catalysts. Acc. Chem. Res. 34, 938–945 (2001).

3 Bell, E. L. et al. Biocatalysis. Nat. Rev. Methods Primers 1, 46 (2021).

4 Wu, S., Snajdrova, R., Moore, J. C., Baldenius, K. & Bornscheuer, U. T. Biocatalysis: Enzymatic Synthesis for Industrial Applications. Angew. Chem. Int. Ed. 60, 88–119 (2021).

5 Arnold, F. H. & Volkov, A. A. Directed evolution of biocatalysts. Curr. Opin. Chem. Biol. 3, 54–59 (1999).

6 Packer, M. S. & Liu, D. R. Methods for the directed evolution of proteins. Nat. Rev. Gen. 16, 379–394 (2015).

7 Zeymer, C. & Hilvert, D. Directed Evolution of Protein Catalysts. Ann. Rev.Biochem. 87, 131–157 (2018).

8 Arnold, F. H. Directed Evolution: Bringing New Chemistry to Life. Angew. Chem. Int. Ed. 57, 4143–4148 (2018).

9 Currin, A. et al. The evolving art of creating genetic diversity: From directed evolution to synthetic biology. Biotech. Adv. 50, 107762 (2021).

10 Xiao, H., Bao, Z. & Zhao, H. High Throughput Screening and Selection Methods for Directed Enzyme Evolution. Ind. Eng. Chem. Res. 54, 4011–4020 (2015).

11 Markel, U. et al. Advances in ultrahigh-throughput screening for directed enzyme evolution. Chem. Soc.Rev. 49, 233–262 (2020).

12 Aharoni, A., Griffiths, A. D. & Tawfik, D. S. High-throughput screens and selections of enzyme-encoding genes. Curr. Opin. Chem. Biol. 9, 210–216 (2005).

13 Tizei, Pedro A. G., Csibra, E., Torres, L. & Pinheiro, Vitor B. Selection platforms for directed evolution in synthetic biology. Biochem. Soc.Trans. 44, 1165–1175 (2016).

14 Taylor, S. V., Kast, P. & Hilvert, D. Investigating and Engineering Enzymes by Genetic Selection. Angew. Chem. Int. Ed. 40, 3310–3335 (2001).

15 Palmer, A. C. & Kishony, R. Understanding, predicting and manipulating the genotypic evolution of antibiotic resistance. Nat. Rev. Gen. 14, 243–248 (2013).

16 Pelletier, J. N., Campbell-Valois, F.-X. & Michnick, S. W. Oligomerization domain-directed reassembly of active dihydrofolate reductase from rationally designed fragments. Proc. Natl. Acad. Sci. U.S.A. 95, 12141–12146 (1998).

17 Rix, G. et al. Scalable continuous evolution for the generation of diverse enzyme variants encompassing promiscuous activities. Nat. Comm. 11, 5644 (2020).

18 Boersma, Y. L. et al. A Novel Genetic Selection System for Improved Enantioselectivity of Bacillus subtilis Lipase A. ChemBioChem 9, 1110–1115 (2008).

19 Reetz, M. T., Höbenreich, H., Soni, P. & Fernández, L. A genetic selection system for evolving enantioselectivity of enzymes. Chem. Comm., 5502-5504 (2008).

20 Kast, P., Asif-Ullah, M., Jiang, N. & Hilvert, D. Exploring the active site of chorismate mutase by combinatorial mutagenesis and selection: the importance of electrostatic catalysis. Proc. Natl. Acad. Sci. U.S.A. 93, 5043–5048 (1996).

21 Yano, T., Oue, S. & Kagamiyama, H. Directed evolution of an aspartate aminotransferase with new substrate specificities. Proc. Natl. Acad. Sci. U.S.A. 95, 5511–5515 (1998).

22 Baker, K. et al. Chemical complementation: A reaction-independent genetic assay for enzyme catalysis. Proc. Natl. Acad. Sci. U.S.A. 99, 16537–16542 (2002).

23 Michener, J. K. & Smolke, C. D. High-throughput enzyme evolution in Saccharomyces cerevisiae using a synthetic RNA switch. Metab. Eng 14, 306–316 (2012).

24 Wang, Q., Tang, S.-Y. & Yang, S. Genetic biosensors for small-molecule products: Design and applications in high-throughput screening. Front. Chem. Sci. Eng. 11, 15–26 (2017).

25 Esvelt, K. M., Carlson, J. C. & Liu, D. R. A system for the continuous directed evolution of biomolecules. Nature 472, 499–503 (2011).

26 Morrison, M. S., Podracky, C. J. & Liu, D. R. The developing toolkit of continuous directed evolution. Nat. Chem. Biol. 16, 610–619 (2020).

27 Wang, L., Brock, A., Herberich, B. & Schultz, P. G. Expanding the Genetic Code of <i>Escherichia coli</i>. Science 292, 498–500 (2001).

28 Chin, J. W. Expanding and reprogramming the genetic code. Nature 550, 53–60 (2017).

29 Young, D. D. & Schultz, P. G. Playing with the Molecules of Life. ACS Chem. Biol. 13, 854–870 (2018).

30 Rubini, R. & Mayer, C. Addicting Escherichia coli to New-to-Nature Reactions. ACS Chem. Biol. 15, 3093–3098 (2020).

31 Martínez-Rodríguez, S., Martínez-Gómez, A. I., Rodríguez-Vico, F., Clemente-Jiménez, J. M. & Las Heras-Vázquez, F. J. Carbamoylases: characteristics and applications in biotechnological processes. App. Microbiol. Biotech. 85, 441–458 (2010).

32 Tack, D. S. et al. Addicting diverse bacteria to a noncanonical amino acid. Nat. Chem. Biol. 12, 138–140 (2016).

33 Clemente-Jiménez, J. M., Martínez-Rodríguez, S., Rodríguez-Vico, F. & Heras-Vázquez, F. J. Optically pure alpha-amino acids production by the “Hydantoinase Process”. Recent Pat. Biotechnol. 2, 35–46 (2008).

34 Altenbuchner, J., Siemann-Herzberg, M. & Syldatk, C. Hydantoinases and related enzymes as biocatalysts for the synthesis of unnatural chiral amino acids. Curr. Opin. Biotech. 12, 559–563 (2001).

35 Martínez-Rodríguez, S., Clemente-Jiménez, J. M., Rodríguez-Vico, F. & Las Heras-Vázquez, F. J. Molecular Cloning and Biochemical Characterization of <i>L</i>-N-Carbamoylase from <i>Sinorhizobium meliloti</i> CECT4114. Microb. Physiol. 9, 16–25 (2005).

36 Martínez-Rodríguez, S. et al. Thermodynamic and mutational studies of l-N-carbamoylase from Sinorhizobium meliloti CECT 4114 catalytic centre. Biochimie 88, 837–847 (2006).

37 Chatterjee, A., Sun, S. B., Furman, J. L., Xiao, H. & Schultz, P. G. A Versatile Platform for Single- and Multiple-Unnatural Amino Acid Mutagenesis in Escherichia coli. Biochemistry 52, 1828–1837 (2013).

38 Jumper, J. et al. Highly accurate protein structure prediction with AlphaFold. Nature 596, 583–589 (2021).

39 Krieger, E. & Vriend, G. YASARA View—molecular graphics for all devices—from smartphones to workstations. Bioinformatics 30, 2981–2982 (2014).

40 Simon, A. J., d’Oelsnitz, S. & Ellington, A. D. Synthetic evolution. Nat. Biotech. 37, 730–743 (2019).

41 Molina, R. S. et al. In vivo hypermutation and continuous evolution. Nat. Rev. Methods Primers 2, 36 (2022).

