## Supplementary Information for "Selecting better biocatalysts by complementing recoded bacteria"

#### Table of contents

1. X-ray crystallography discussion S2
2. Methods S4
3. Crystallography and structure determination S16
4. Sequences S18
5. Supporting Figures S20
6. NMR spectra S21
7. References S23

#### 1. X-ray crystallography discussion

**General structural features:** Monomeric SmLcar (Gly5 – Glu416) variants fold into a catalytic domain, (residues 5-215 and 330-416) that comprises the metal binding sites and the active site, and a dimerization domain (residues 216-329). The individual domains are connected by a linker region that is referred to as the "hinge region". The obtained structures are comparable to other members of the amidase, hydantoinase/carbamoylase family (InterPro accession IPR010158). Ligand-induced conformational changes are well characterized in this family, in which the catalytic domain can rotate up to 32° with respect to the dimerization domain, resulting in an open or closed conformation. In the closed conformation some active residues are closer to the substrate.[1,2] Here, the two determined SmLcar structures are in the closed state, suggesting they crystallized in a catalytically active state.

**Metal-binding site:** The catalytic domain of SmLcar variants contains a bi-metal center in the active site. In the homologous structures these metals are modeled as $Zn^{2+}$ or $Mn^{2+}$ atoms, although supplementing carbamoylases with other bivalent metals such as $Co^{2+}$, $Ni^{2+}$, or $Fe^{2+}$ also results in catalytically active assemblies.[3,4] Crystals of SmLcar variants were grown without addition of metal ions, which makes it *a priori* impossible to determine the nature of them. In both monomers of the SmLcar and SmLcar_GC structures (measured at 0.965 Å), two anomalous peaks were observed at the putative metal sites, indicating Mn ($f$'=1.3 e), Zn (2.5 e), Ni (1.9 e), Co (1.7 e) or Fe (1.5 e). To elucidate the nature of the metal ion, we determined the metal content of purified SmLcar variants by ICP-MS. While for SmLcar (monomer concentration = 100 µM), this analysis gave 60 µM Zn and 90 µM Fe, for SmLcar_GC (600 µM) yielded 540 µM of Zn and 200 µM of Fe. For the latter analysis, Mn, Ni and Co were present at concentrations lower than 30 µM. As ICP-MS is not able to differentiate which metal is bound at which site, we refined the crystal structures with partial occupancies, that is all metal sites containing both $Fe^{2+}$ and $Zn^{2+}$. Occupancies used were 0.40 for $Zn^{2+}$ and 0.34 for $Fe^{2+}$ in SmLcar and $0.44^{2+}$ for Zn and 0.16 for $Fe^{2+}$ in SmLcar_GC. The interacting residues His87, Asp98 (bidentate), His194 and His386 coordinate the ions in a trigonal bipyramidal fashion with geometry and ligation distances being consistent with $Zn^{2+}$ and $Fe^{2+}$ ions, as confirmed using the *CheckMyMetal* web server.[5]

**Substrate binding site:** The substrate binding site is located adjacent to the bi-metal center at the interface of the catalytic domain of one monomer and the dimerization domain of the other. In their closed conformations, the binding site of SmLcar_GC (82 $Å^3$) is significantly larger than the one of the wild-type (16 $Å^3$). In SmLcar Leu217 and Phe329 are located next to Arg292 and have hydrophobic interaction with each other, while in SmLcar_GC the active site is opened up by the introductions of the smaller amino acids, glycine and cysteine, respectively.

While no ligands were added during protein crystallization, we identified electron density corresponding to small-molecules in the crystal structures of SmLcar and SmLcar_GC. We tentatively assigned these as ornithine in SmLcar and carbamoyl-*L*-tyrosine in SmLcar_GC, with both molecules making extensive interactions with adjacent side chains. For example, the conserved Arg292 is making a salt bridge with the carboxyl moiety of ornithine in the SmLcar structure. Furthermore, several hydrogen bonds between amino acid side chains (e.g. Glu132, His230, Asn279 or Gly36) and ornithine can be discerned.

#### 2. Methods

**Materials & Methods:** Chemicals, including *L*-3nY and *L*-3iY, were purchased from *Sigma Aldrich* and used without further purification unless otherwise noted. $^{1}$H-NMR and $^{13}$C-NMR spectra were recorded on a Bruker 400 in DMSO-*d6*. Plasmid pUC57-3iYaaRS, bearing the gene encoding the 3iY aminoacyl tRNA synthetase (3iYaaRS)[6], the 2MCS synthetic gBlock fragment and the SmLcar coding sequence were purchased from *GenScript* (USA); pACYCDuet-1 was purchased from *MilliporeSigma* (Novagen), pULTRA-CNF was a gift from Prof. Dr. Peter Schultz (Addgene plasmid #48215). *Escherichia coli* strains NEB10-beta and BL21(DE3) (*New England Biolabs*) were used for cloning and expression experiments. Standard primers were synthesized by *Eurofins MWG Operon* (Germany), while degenerate primers by *Biolegio* (The Netherlands). Plasmid Purification Kits were obtained from *QIAGEN* (Germany) and DNA sequencing carried out by *Eurofins* (Germany). Phusion polymerase, T4 ligase, *Esp3I* and *BsaI* were purchased from *New England Biolabs*; Ni-NTA resin (Ni Sepharose$^{TM}$ 6 Fast Flow) from *GE Healthcare Life Sciences* (Germany).

The concentration of DNA in solutions was determined based on the absorption at 260 nm on a Thermo Scientific Nanodrop 2000 UV-Vis spectrophotometer. Cellular density ($OD_{600}$) was measured on an Ultrospec 10 Cell Density Meter (Biochrom). Analytical HPLC analysis was performed on a Waters Acquity HPLC class system (Waters) equipped with a PDA detector. All analyses were performed using a reverse-phase HPLC column (XSelect-CSH-$C_{18}$, 5 µm, 4.5×150 mm; Waters) kept at 40 °C and the sample plate was kept at room temperature. Samples were separated with a gradient from 5 to 95% acetonitrile (0.1% TFA) in water (0.1% TFA) at a flow rate of 1.0 mL/min. Absorbance was monitored at different wavelengths for the different ncAAs and their carbamoylated derivatives ($\lambda$ = 359 nm for 3nY, $\lambda$ = 282 nm for 3iY). Each ncAA or cam-ncAA was quantified according to a calibration curve obtained from samples containing different concentrations of authentic standards (**Fig. S1**).

**Synthesis of cam-3iY and cam-3nY:** Carbamoylated ncAA precusors were synthesized following a report by Lenman *et al.* with minor modifications.[7] In brief, to a suspension of the ncAA (2 mmol, 1 eq) in water (20 mL), KOCN (22 mmol, 11 eq.) was added. The reaction was stirred for 4 hours at 60 °C under $N_2$ atmosphere. The reaction mixture was placed in an ice bath and acidified to pH 1 with 2 M HCl. The resulting precipitate was filtered off and extensively washed with ice-cold water and subsequently dried under vacuum to yield carbamoylated ncAAs in good purity and moderate yields.

**cam-3iY:** white solid, 51% yield: **$^1$H NMR** (400 MHz, DMSO-*d6*) δ 10.11 (s, 1H), 7.44 (d, J = 2.1 Hz, 1H), 6.97 (dd, J = 8.2, 2.1 Hz, 1H), 6.76 (d, J = 8.2 Hz, 1H), 6.09 (d, J = 8.2 Hz, 1H), 5.60 (s, 2H), 4.22 (td, J = 7.8, 5.2 Hz, 1H), 2.84 (dd, J = 13.9, 5.2 Hz, 1H), 2.70 (dd, J = 13.8, 7.6 Hz, 1H). **$^{13}$C NMR** (101 MHz, DMSO-d6) δ 174.31, 158.54, 155.64, 139.62, 130.76, 130.45, 115.09, 84.73, 54.26, 36.59.

**cam-3nY:** yellow solid, 36% yield. **$^1$H NMR** (400 MHz, DMSO-*d6*) δ 10.83 (s, 1H), 7.68 (d, J = 2.2 Hz, 1H), 7.36 (dd, J = 8.5, 2.2 Hz, 1H), 7.05 (d, J = 8.5 Hz, 1H), 6.19 (d, J = 8.3 Hz, 1H), 5.61 (s, 2H), 4.30 (td, J = 8.0, 5.1 Hz, 1H), 2.98 (dd, J = 13.9, 5.2 Hz, 1H), 2.83 (dd, J = 13.9, 7.9 Hz, 1H). **$^{13}$C NMR** (101 MHz, DMSO-d6) δ 174.13, 158.49, 151.37, 136.77, 136.63, 129.18, 125.82, 119.38, 53.96, 36.68.

**Construction of pACYC_GG:** In order to streamline the cloning efforts, a plasmid enabling the modular exchange of the target enzyme and/or the β-lactamase via Golden Gate assembly was constructed.[8] This *pACYC_GG* plasmid is based on the commercially available pACYCDuet-1, a bacterial vector with a p15A origin of replication and featuring two coding sequences. We redesigned both multiple cloning sites (MCSs) in order to be able to insert DNA fragments using two different type IIS restriction enzymes: *BsaI* and *Esp3I.* To achieve this, we first suppressed the three *Esp3I* recognition sites that were present in the original plasmid backbone. For this, we made use of three pairs of primers (*pACYCGG_1, pACYCGG_2, pACYCGG_3, for* and *rev,* see *Sequences*) to insert silent mutations and to obtain the modified plasmid backbone pACYC_Sup by Gibson assembly of the three resulting DNA fragments. Successful assembly and mutagenesis were confirmed by sequencing with *Esp3ISup_Check1* and *_Check2*. Next, the redesigned MCSs were synthesized as one gBlock Gene Fragment (2MCS) with strategically placed *BsaI* and *Esp3I* sites for Golden Gate assembly in the first and second MCS, respectively. To generate the final pACYC_GG plasmid, another Gibson assembly was performed between: (1) the aforementioned 2MCS gBlock and (2) the linearized modified backbone obtained via PCR using *pACYClin_for* and *_rev* primers on pACYC_Sup. The assembly reaction was used to transform chemically competent *E. coli* NEB10-beta cells. A single colony was picked from LB plates containing chloramphenicol (35 µg/mL) and used to inoculate 5 mL of LB medium containing the same concentration of chloramphenicol. Bacteria were grown over night, plasmids isolated and successful assembly of pACYC_GG verified by sequencing with *DuetUP1*.

**Assembly of the selection plasmid:** PCR products and/or synthetic genes flanked by the appropriate type IIS recognition sites can be easily assembled into either MCS1 (using *BsaI*) or MCS2 (using *Esp3I*) of pACYC_GG, either individually or in parallel in a one-pot reaction with T4 DNA ligase together with *BsaI* and/or *Esp3I.* The following thermocycler program was followed for the assemblies: (1) 50 cycles alternating between 37 ˚C and 16 ˚C for 5 and 10 min respectively, (2) a final digestion step at 50 ˚C for 20 min, and (3) an enzyme inactivation step at 80 ˚C for 10 min. The restriction-ligation reactions were transformed into chemically competent *E. coli* NEB10-beta cells. A single colony was picked from LB plates containing chloramphenicol (50 µg/mL) and used to inoculate 5 mL of LB medium containing the same concentration of chloramphenicol. Bacteria were grown over night, plasmids isolated and successful assembly verified by sequencing with either *DuetUP1* and *DuetDOWN1* for MCS1-cloning or *DuetUP2* and *T7 Terminator* for MCS2-cloning. Carbamoylases were cloned into MCS1, TEM-1.B9[9] was cloned into MCS2. For this, the synthetic gene coding for SmLcar flanked by BsaI sites (see Sequences section) could be directly assembled in pACYC_GG as previously described, while the CDSs of the engineered variants were first amplified via PCR from the correspondent pACYC plasmids using SmLcarCDS_GG_for and _rev primers and subsequently assembled. The CDS of TEM-1.B9 flanked by Esp3I sites was obtained via PCR starting from the plasmid pAddict[10] using the TEM1.B9_GG_for and _rev primers before assembly.

**Construction of pULTRA_3iY:** The plasmid bearing the 3iY aminoacyl tRNA synthetase was designed based on the pULTRA-CNF vector by simply swapping the two CDSs, while retaining the same tRNA cassette. The CDS of 3iYaaRS was amplified by PCR using *3IYRS_CDS_for* and *_rev* primers on pUC57-3iYaaRS as template, while the pULTRA backbone was linearized using *pULTRA_lin_for* and *_rev*. The resulting DNA fragments were purified and assembled by Gibson, before transforming chemically competent *E. coli* NEB10-beta cells. A single colony was picked from LB plates containing spectinomycin (50 µg/mL) and used to inoculate 5 mL of LB medium containing the same concentration of spectinomycin. Bacteria were grown over night, plasmids isolated and successful assembly of pULTRA_3iY verified by sequencing with *3IYRS_CDS_for* and *_rev*.

**Assembly of the chemical complementation platform:** pULTRA_3iY plasmid was transformed into chemically competent *E. coli* BL21(DE3) and a single colony was used to inoculate an overnight culture. Chemically competent *E. coli* BL21(DE3) bearing pULTRA_3iY were prepared and subsequently co-transformed with a selection plasmid featuring the desired carbamoylase variants. A single colony was picked from LB plates containing spectinomycin (50 µg/mL) and chloramphenicol (35 µg/mL) and used to inoculate 5 mL of LB medium containing the same antibiotics.

**Site-directed mutagenesis:** Starting from pACYC_SmLcar, mutagenic primers *SmLcar_KO_for* and *_rev* were used to generate a knocked-out (KO) variant of SmLcar with the Arg292Ala substitution. The following PCR protocol was used: (1) initial denaturation 95 ˚C for 3 min, (2) 16 cycles of denaturation at 95 ˚C for 30 s, annealing at 58 ˚C for 30 s and extension at 72 ˚C for 1 min 30 s; (3) a final extension at 72 ˚C for 10 min. The resulting PCR product was digested with *DpnI* for 1 hour at 37 ˚C, purified, and transformed into chemically competent *E. coli* NEB10-beta cells. A single colony was picked from LB plates containing chloramphenicol (35 µg/mL) and used to inoculate 5 mL of LB medium containing the same concentration of chloramphenicol. Bacteria were grown over night, plasmids isolated and variants harboring the correct mutations identified by sequencing.

After the second round of directed evolution, one of the best variants identified from the Leu217Gly_Phe329NNK library was also bearing an unexpected point mutation (G to A) in a position that was not targeted, resulting in SmLcar_Leu217Gly_Phe329Tyr_Ala346Thr. Intriguingly, this mutation was not detected in any other hit that we sequenced and it is far from the catalytic pocket. To test whether the Ala346Thr substitution has any effect on the catalytic performance of SmLcar_GY, we reverted the mutation as described above by using the mutagenic primers *T346A_for* and *_rev*. As expected, this substitution did not result in a significant change in catalytic activity (**Fig. S3**).

**Validation of the chemical complementation platform in 96-well plates:** LB medium (6 mL) containing 25 µg/mL spectinomycin and 17.5 µg/mL chloramphenicol were inoculated with 50 µL of a densely grown overnight culture of *E. coli* BL21(DE3) cells harboring pULTRA-3iY and the appropriate selection plasmid. Cells were incubated at 37 °C and 135 rpm until an optical density at 600 nm ($OD_{600}$) of 0.4–0.6 was reached. At this point, gene expression was induced with IPTG (final concentration 1 mM), and cells were incubated at 37 °C and 135 rpm for 3 more hours. In parallel, 5 mL of LB medium with 25 µg/mL spectinomycin, 17.5 µg/mL chloramphenicol, 0.1 mM IPTG and increasing concentrations of carbenicillin (from 0 µg/mL to 150 µg/mL) was freshly prepared. Induced cultures were then all diluted to an $OD_{600}$ of 0.2 using LB medium with 25 µg/mL spectinomycin, 17.5 µg/mL chloramphenicol, 0.1 mM IPTG. Diluted cultures and LB medium containing increasing concentrations of carbenicillin were transferred to a transparent 96-well assay plate that was set up by adding in each well: (1) 135 µL of LB containing the carbenicillin concentration gradients (final concentrations ranging from 0 µg/mL to 100 µg/mL), (2) 45 µL of the diluted cultures, (3) 10 µL of cam-3nY/3nY stock solution (both 10 mM in LB medium), and (4) 10 µL of $MnCl_2$ (2.5 mM in 50 mM $Na_2HPO_4$, 150 mM NaCl, pH 8, final concentration 125 µM). At this stage, the assay plate was transferred into a Synergy H1 microplate reader (*BioTek*) that was preheated to 30 °C. Optical/cell density at 600 nm ($OD_{600}$) was measured every 10 minutes from the bottom of the wells for 36 hours while continuously shaking (double orbital, 425 c.p.m.).

***Alphafold* model of SmLcar and residue selection mutagenesis:** To identify positions that, when targeted, could influence the activity of SmLcar for ncAA precursors, the threedimensional structure of SmLcar was predicted using ColabFold[11] and assuming a homo-dimeric organization. Residues for randomization were chosen after visual inspection of the model based on the following two criteria: (1) multiple sequence alignment of predicted/known carbamoylases identified the residues as non-conserved and (2) residues are in proximity to the catalytic cavity of the aforementioned homology model without being essential for enzymatic activity.[12] Ultimately, four residues (Gln91, Val150, Leu217, and Phe329) were selected following cross-validation of candidates with residues suggested by the online tool HotSpot Wizard 3.0.[13]

**Generation of NNK libraries:** Overlap extension PCR (oePCR) with primers bearing degenerate NNK codons was employed to randomize the previously identified positions. Starting from pACYC_SmLcar, four different *NNK_for* mutagenic primers (Gln91, Val150, Leu217, or Phe329) were used in combination with *SmLcarCDS_GG_rev* to generate four PCR fragments. Partially overlapped PCR fragments were obtained using the corresponding *_rev* primer (again specific for each of the four positions) in combination with *SmLcarCDS_GG_for*. In the final step, each fragment was coupled with the equivalent partially-overlapping one and the full-length, 1280 bp oePCR products were obtained by running the reactions in presence of both *SmLcarCDS_GG_for* and *_rev* primers. The following PCR protocol was used: (1) initial denaturation at 95 ˚C for 3 min, (2) 30 cycles of denaturation at 95 ˚C for 30 s, annealing at 60 ˚C for 30 s and extension at 72 ˚C for 20 s; (3) a final extension at 72 ˚C for 10 min. After each PCR, the fragment of interest was isolated and purified via agarose gel electrophoresis. Finally, the partially-randomized CDS was cloned in pACYC_TEM-1.B9 using *BsaI* as previously described. The restriction-ligation reactions were transformed into chemically competent *E. coli* NEB10-beta cells first. All colonies were scraped from LB plates containing chloramphenicol (35 μg/mL) and plasmid DNA was extracted, isolated and library quality verified by sequencing with *DuetUP1* and *DuetDOWN1*. Finally, 300-400 ng of the obtained plasmid DNAs were transformed into chemically competent *E. coli* BL21(DE3) already bearing pULTRA_3iY and spread onto LB agar plates containing spectinomycin (50 μg/mL) and chloramphenicol (35 μg/mL). After incubating overnight at 37 °C single colonies were picked and transferred into 96-deep well plates containing LB media with the same antibiotics. In the second round of directed evolution, the oePCR was performed with the coding sequence of SmLcar_G as a template, targeting the remaining three positions.

**Chemical complementation screening in liquid media:** As previously described, 96-deep well plates filled with LB medium (0.5 mL) containing 50 μg/mL spectinomycin and 35 μg/mL chloramphenicol were inoculated with a single colony of *E. coli* BL21(DE3) cells harboring pULTRA-3iY and one version of mutagenized CDS cloned into pACYC_TEM-1.B9. In addition to library members, single colonies of freshly transformed controls (SmLcar, SmLcar_KO, and SmLcar_L217G in the second round, all cloned into pACYC_TEM-1.B9) were used as controls. The resulting 96-deep well plates were incubated overnight at 37 °C while shaking at 900 rpm (*Titramax 1000 & Incubator 1000, Heidolph*). The next morning, 10 μL of the densely grown overnight cultures were transferred into fresh 96-deep well plates containing 0.5 mL LB media and appropriate antibiotics. Bacteria were cultured at 37 °C for 1.5 hours while shaking at 900 rpm. Subsequently, protein production was induced by addition of 16.5 μL of 30 mM IPTG (final concentration of 1 mM), and plates were incubated at 37 °C for 3 more hours while shaking (900 rpm). Induced cultures were then all diluted to an $OD_{600}$ of 1 using LB medium with 25 μg/mL spectinomycin, 17.5 μg/mL chloramphenicol, and 0.1 mM IPTG. Diluted cultures and LB media containing 25 μg/mL spectinomycin, 17.5 μg/mL chloramphenicol, 0.1 mM IPTG, and a fixed concentration of carbenicillin were transferred to a transparent 96-well assay plate that was set up by adding in each well: (1) 170 μL of LB with either 118 or 235 μg/mL of carbenicillin (final concentration 100 or 200 μg/mL, respectively), (2) 10 μL of diluted culture, (3) 10 μL of 10 mM cam-3nY stocks in LB (final concentration 500 μM) or 10 mM 3nY stocks in LB (final concentration 500 μM), and (4) 10 μL of 2.5 mM $MnCl_2$ in 50 mM $Na_2HPO_4$, 150 mM NaCl, pH 8 (final concentration 125 μM). At this stage, the assay plate was transferred into a Synergy H1 microplate reader (*BioTek*) that was preheated to 30 °C. Optical/cell density at 600 nm ($OD_{600}$) was measured every 10 minutes from the bottom of the wells for 36 hours while continuously shaking (double orbital, 425 c.p.m.).

**Production and purification of CLs**: Flasks containing 500 mL LB medium with 35 μg/mL chloramphenicol were inoculated with 500 μL of a densely grown overnight culture of *E. coli* BL21(DE3) cells harboring the appropriate pACYC_CL plasmid. Cells were incubated at 37 °C and 135 rpm until an optical density at 600 nm of around 0.6 was reached and gene expression was induced by adding IPTG (final concentration 0.1 mM). Enzymes were produced for 4 hours at 37 °C while shaking (135 rpm), after which cells were harvested by centrifugation (3,700 rpm for 10 min, 4 °C). Cell pellets were resuspended in buffer (20 mL, 50 mM $Na_2HPO_4$, 500 mM NaCl, pH 8, containing 1 mg/mL egg white lysozyme and half a tablet of protease inhibitor cocktail (*Roche*). The cells were then lysed by sonication (10 min, 5 s pulse and pause cycles, 70% amplitude)**,** and cellular debris was removed by centrifugation (12,000 rpm for 45 min, 4 °C). The supernatant was loaded onto a Ni-NTA resin and purified according to the manufacturer's specifications. Protein-containing fractions were pooled after elution, concentrated, and finally stored in 50 mM $Na_2HPO_4$, 150 mM NaCl, pH 7 with 5% glycerol at -20 °C. The identity of CLs was confirmed by SDS-PAGE (**Fig. S2)** and mass spectrometry (Q-TOF MS, see Table below).

| Carbamoylase variant | Expected (M-Met, Da**)** | Found (M, Da) |
|---|---|---|
| **SmLcar** | 46850.13 | 46857 |
| **SmLcar_G** | 46794.02 | 46801 |
| **SmLcar_GC** | 46749.98 | 46756 |
| **SmLcar_GY** | 46840.05 | 46846 |
| **SmLcar_GY – A346T** | 46810.02 | 46804 |

**Size exclusion chromatography (SEC):** Before protein crystallization trials, another chromatographic step was performed in order to exchange the buffer. Molecular mass of native enzymes and their oligomeric state were also determined by size exclusion chromatography. Superdex 10/300, 200 increase (*Cytiva*) column was equilibrated with 50 mM Tris-HCl, 150 mM NaCl, pH 7.5. The flow rate for protein elution was 0.3 mL $min^{-1}$. Thyroglobulin (MW 670 kDa), γ-globulin (MW 158 kDa), Ovalbumine (MW 44 kDa), Myoglobin (MW 17 kDa) and Vitamin B12 (MW 1350 Da) were used as reference standards for the calibration curve. SmLcar and its variants displayed an estimated native MW around 75 kDa, suggesting a homodimeric structure in solution.

**In vitro characterization of carbamoylases:** Standard enzymatic reactions were carried out with the purified protein (at a final concentration of either 10 µM for SmLcar or 0.5 µM for the evolved variants) dissolved in 50 mM $Na_2HPO_4$, 150 mM NaCl in 200 µL reaction volume. The pH of the buffer was different depending on the substrate: either 8.0 (with 2 mM of cam-3iY) or 6.0 (2 mM cam-3nY). The reaction mixtures were incubated at room temperature and aliquots were quenched by addition of one volume equivalent of 3% $H_3PO_4$. After centrifugation, the resulting supernatants were analyzed by HPLC. The concentrations of cam-ncAAs and ncAAs were determined as described in the Materials & Methods section **(Fig. S1)**.

**Kinetic characterization of SmLcar variants:** Steady-state kinetics studies in presence of different (0.6-20 mM) cam-3nY/cam-3iY concentrations were performed at room temperature in 100 mM $Na_2HPO_4$, 100 mM NaCl, 0.5 mM $MnCl_2$, at pH 6 or 8. Reactions were quenched at varying time points for different SmLcar variants by addition of one volume equivalent of

3% $H_3PO_4$ (see Table below). After centrifugation, the resulting supernatants were analyzed by HPLC as described in the previous section.

| Carbamoylase | cam-3nY | | | cam-3iY | | |
|---|---|---|---|---|---|---|
| | [E] (µM) | pH | Time (min) | [E] (µM) | pH | Time (min) |
| **SmLcar** | 6 | 6 | 1440 | 10 | 8 | 1440 |
| **SmLcar_G** | 1 | 6 | 60 | 1 | 8 | 60 |
| **SmLcar_GY** | 1 | 6 | 18 | 1 | 8 | 60 |
| **SmLcar_GC** | n.d. | n.d. | n.d. | 1 | 6 | 30 |

**Mock selections:** Precultures of 5 mL LB media containing 50 µg/mL spectinomycin and 35 µg/ml chloramphenicol were inoculated from glycerol stocks of *E. coli* BL21(DE3) cells harboring pULTRA-3iY and the selection plasmid featuring either SmLcar or SmLcar_GY. Following overnight growth at 37 °C while shaking at 135 rpm, cultures with mixed populations were started by combining the densely grown precultures in appropriate amounts as follows: 25 µL of SmLcar cells and 25 µL of SmLcar_GY cells (1:1), or 50 µl of SmLcar cells and 50 µl of a ten times-diluted sample of SmLcar_GY cells (10:1) were added to 5 mL LB media containing 25 µg/mL spectinomycin and 17.5 µg/mL chloramphenicol. The resulting populations were cultured at 37 °C and 135 rpm until an $OD_{600}$ of 0.3-0.5 was reached. At this point, gene expression was induced with IPTG (final concentration 0.1 mM) and cells were incubated at 37 °C and 135 rpm for 3-4 more hours, until $OD_{600}$ ~1. During this step, selection medium was freshly prepared containing 25 µg/mL spectinomycin, 17.5 µg/mL chloramphenicol, 0.1 mM IPTG, 125 µM $MnCl_2$, 500 µM cam-3nY, and 50 µg/mL carbenicillin. Positive control medium was prepared in the same manner, except with 500 µM 3nY instead of cam-3nY. Induced cultures were then diluted 1:100 in 5 mL selection medium and grown for 24-48 hours at 30 °C and 135 rpm, while routinely measuring $OD_{600}$. Upon reaching $OD_{600}$ > 0.8, serial dilutions (1:100) were performed in fresh selection medium. For each selection campaign, two to three rounds of selection were performed as described until the genotype of the population had converged. Prior to selection and after each serial passage, overnight cultures of the mixed populations were inoculated in LB with 50 µg/mL spectinomycin and 35 µg/ml chloramphenicol. Plasmids were isolated and variants in the population were identified by Sanger sequencing with *DuetDOWN1*. Base calls taken from raw sequencing data were used to calculate the relative distribution of bases at positions of interest.

**Selections of improved SmLcar variants from libraries:** Chemically competent *E. coli* BL21(DE3) bearing pULTRA_3iY were transformed with selections plasmids containing either the SmLcar_Leu217NNK or SmLcar_G_F329NNK library. After recovery, transformants were not plated but instead grown as a preculture in 4 mL SOC medium containing 50 µg/mL spectinomycin and 35 µg/ml chloramphenicol. The next morning, cultures of 5 mL LB containing 25 µg/mL spectinomycin and 17.5 µg/mL chloramphenicol were started with 50 µL of the densely grown overnight culture. Selections were performed as described above with exception of using 20 µg/mL carbenicillin for the first growth cycle of the SmLcar_Leu217NNK library. After three growth-dilution cycles for the SmLcar_G_F329NNK library, cells were plated on LB agar containing 50 µg/mL spectinomycin and 35 µg/ml chloramphenicol and grown overnight at 37 °C. Ten single colonies were picked and used to inoculate 4 mL of LB medium containing the same antibiotics. Bacteria were grown overnight, plasmids isolated and the SmLcar variant identified by sequencing with *DuetDOWN1*.

#### 3. Crystallization and structure determination

SmLcar and SmLcar_GC (SmLcar_L217G _F329C) were produced and purified as described above. Purified proteins were concentrated to ~10 mg/mL in 50 mM Tris buffer (pH 7.5) containing 150 mM NaCl. SmLcar protein crystals were grown from 0.2 M sodium acetate, 0.1 M Bis-tris propane pH 6.5 and 20% PEG3350. SmLcar_GC protein crystals were grown from 20 mM Na,K phosphate, 0.1 M Bis-tris propane pH 6.5, 20% PEG3350. Prior to flash-cooling in liquid $N_2$, crystals were cryoprotected with 25% glycerol supplemented to the mother liquor. X-ray diffraction data for SmLcar and SmLcar_GC were collected at ESRF beamline MASSIF-1.[14] Diffraction data were (re-)processed with the program XDS[15] and were (re-)indexed with AIMLESS[16] from CCP4[17]. Molecular replacement was performed with PHASER[18], using the aforementioned *AlphaFold2* models with two domains as search models. The SmLcar and SmLcar_GC crystals belong to the monoclinic space group *I2* with two monomers of 47 kDa in the asymmetric unit. The $V_M$ is 2.0 Å$^3$/Da with a solvent content of 38%[19]. The construct contains a 21 residue N-terminal extension, including a His-tag that is not observed in electron density. The resulting structures were improved by several rounds of model building and refinement, using the programs Coot[20] and REFMAC5[21]. Metal sites were evaluated with the *CheckMyMetal* webserver[22]. The quality of the models were analyzed with PDB_REDO[23] and MolProbity[24]. The CastP server[25] with default values was used for calculating the volume of the active sites (http://sts.bioe.uic.edu/castp/calculation.html). Polder omit maps (Liebschner, 2017) were calculated with *Phenix*.[26]

**Table 1.** Data collection and refinement statistics. Values in parentheses are for the highest resolution shell.

| Diffraction data | SmLcar | SmLcar_GC |
|---|---|---|
| Wavelength (Å) | 0.965 | 0.965 |
| Resolution range (Å) | 45.5 – 1.75 | 99.2 -2.30 |
| Space group | *I2* | *I2* |
| Cell dimensions (Å) *a, b*, *c, β* | 133.3, 41.8, 134.0, 94.1 | 132.6, 42.1, 137.2, 94.8 |
| Unique reflections | 74615 (3970) | 33664 (2938) |
| Completeness (%) | 99.5 (97.4) | 97.7 (89.8) |
| Overall I/σ (I) | 13.3 (1.9) | 9.9 (1.2) |
| $R_{merge}$ (%) | 8.0 (70.8) | 10.6 (91.4) |
| $R_{pim}$ (%) | 3.7 (33.2) | 5.2 (49.7) |
| *R/ Rfree* (%) | 14.9 / 18.6 | 17.3 / 23.8 |
| r.m.s. deviations from ideal values | | |
| Bond lengths (Å) | 0.010 | 0.012 |
| Bond angles (°) | 1.41 | 1.50 |
| Protein residues | 822 | 820 |
| Metal atoms<br>Occupancy | 4 x ($Zn^{2+}$, $Fe^{2+}$)<br>0.40, 0.34 | 4 x ($Zn^{2+}$, $Fe^{2+}$)<br>0.44, 0.16 |
| Ligand molecules | 2 x ornithine | 2 x cam-*L*-tyrosine |
| Water molecules | 466 | 135 |
| Ramachandran favored (%) | 97.8 | 98.0 |
| Ramachandran allowed (%) | 2.2 | 2.0 |
| Ramachandran outliers (%) | 0.0 | 0.0 |
| Clashscore | 3.38 | 2.04 |
| MolProbity score | 1.17 | 1.09 |
| PDB accession ID | 8APZ | 8AQ0 |

#### 4. Sequences

Sequences of primers with mutations / randomizations shown in bold

| Name | Sequence 5'→3' |
|---|---|
| pACYClin_rev | GTATATCTCCTTATTAAAGTTAAAC |
| pACYClin_for | AAAACCTAGGCTGCTGCCACC |
| DuetUP2 | TTGTACACGGCCGCATAATC |
| DuetDOWN1 | GATTATGCGGCCGTGTACAA |
| DuetUP1 | GGATCTCGACGCTCTCCCT |
| T7 Terminator | GCTAGTTATTGCTCAGCGG |
| pACYCGG_2for | CACCCAGGGATTGGCTGAAACGAAAAACATATTCTCAA |
| pACYCGG_1rev | TTGAGAATATGTTTTTCGTTTCAGCCAATCCCTGGGTG |
| pACYCGG_3for | TACGTGCCGATCAACCTCTCATTTTCGCCAAAAGTT |
| pACYCGG_2rev | AACTTTTGGCGAAAATGAGAGGTTGATCGGCACGTA |
| Esp3ISup_Check1 | ATACACTAAATCAGTAAGTTGGC |
| Esp3ISup_Check2 | CACCGGAAGGAGCTGACTGG |
| pACYCGG_1for | GTTTTTCTTTTCACCAGTGAAACGGGCAACAGCTGATTGCCC |
| pACYCGG_3rev | GGGCAATCAGCTGTTGCCCGTTTCACTGGTGAAAAGAAAAAC |
| 3IYRS_CDS_for | ATGGACGAATTTGAAATGATAAAGAG |
| 3IYRS_CDS_rev | TTATAATCTCTTTCTAATTGGCTCTAAAATC |
| pULTRA_lin_for | GATTTTAGAGCCAATTAGAAAGAGATTATAAGCGGCCGCGTTTAA ACGGTC |
| pULTRA_lin_rev | CTCTTTATCATTTCAAATTCGTCCATGCGGCCGCACCTCCTTTGT GA |
| SmLcar_KO_for | CACAGTAGACATC**GCC**TCCCCTGATCAAGCAAAGC |
| SmLcar_KO_rev | ATCAGGGGA**GGC**GATGTCTACTGTGAAAACTACTT |
| T346A_for | GTTCGTGGGGCG**G**CAGAGAAATTAGGGTATTCGC |
| T346A_rev | TTCTCTG**C**CGCCCCACGAACTGTCTCTACAAG |
| Q91NNK_for | CTCGCACCTGGATACT**NNK**CCTACCGGAGGAAAATTTGACGG |
| Q91_rev | AGTATCCAGGTGCGAGCCAATATGAAC |
| V150NNK_for | TGGCGTATTTGCAGGC**NNK**CACACCTTAGAATATGCCTACG |
| V150K_rev | GCCTGCAAATACGCCAGAGGCCAACATTGC |
| L217NNK_for | TCACTCATTGTCAAGGC**NNK**TGGTGGCTGGAATTTACTTTGAC |
| L217_rev | GCCTTGACAATGAGTGACAACGCCAATTTG |
| F329NNK_for | ATTGAAGCAGTAGGTCAC**NNK**GACCCAGTAACCTTTGACCCT |
| F329_rev | GTGACCTACTGCTTCAATCGAGCATCCGACAC |
| K369NNK_for | CGCCTGCTGGGCGGCT**NNK**GTTGCACCTACGACAATGATCATG |
| K369_rev | AGCCGCCCAGCAGGCGTCGTGTC |
| SmLcarCDS_GG_fo r | ACCGGTCTCCTGGTATGGCGGCGCCTGGT |
| SmLcarCDS_GG_re v | GTTGGTCTCCCAAGTCACTCCACAATCTC |
| TEM1.B9_GG_for | GCATCGTCTCATCGGCAGCATGAGTATTCAACATTTCC |
| TEM1.B9_GG_rev | ATGCCGTCTCAGGTCGCTGCCTTACCAATGCTTAATCAG |

**Sequence 5’→3’ of 2MCS synthetic gBlock:**

CCTGTAGAAATAATTTTGTTTAACTTTAATAAGGAGATATACCATGGGCAGCAGCCATCATC
ATCATCATCACGGCAGCGGCCTGGTGCCGCGCGGCAGCGCTGGTAGAGACCGGGCCTGAAGG
TCTCGCTTGGGCCCGAACAAAAACTCATCTCACGAACAGAAAGTAATCGTATTGTACACGGC
CGCATAATCGAAATTAATACGACTCACTATAGGGGAATTGTGAGCGGATAACAATTCCCCAT
CTTAGTATATTAGTTAAGTATAAGAAGGAGATATACATATGTTCGGCGAGACGGAAAGTGAA
ACGTGATTTCATGCGTCATTTTGAACATTTTGTAAATCTTATTTAATAATGTGTGCGGCAAT
TCATGCGTTTATACGTCTCTGACCGGAAAGAAACCGCTGCTGCGAAATTTGAACGCCAGCAC
ATGGACTCGTCTACTAGCGCAGCTTAATTAACCTAGGAGAAATAAAACCTAGGCTGCTGCCA
CCGCTGAGCAATAACTAGC

**Sequence 5’→3’ of SmLcar synthetic gene (CDS flanked by *BsaI* sites):**

CTAACCGGTCTCCTGGTATGGCGGCGCCTGGTGAAAACCGTCGTGTAAATGCCGATCGCTTA
TGGGACTCGCTGATGGAAATGGCGAAAATTGGCCCTGGAGTTGCGGGAGGTAACAACCGCCA
GACCTTGACAGACGCTGACGGGGAAGGGCGCCGCCTGTTTCAATCCTGGTGTGAAGAAGCCG
GCTTGTCGATGGGCGTCGATAAGATGGGCACCATGTTTCTTACACGCCCTGGAACTGACCCC
GACGCATTGCCAGTTCATATTGGCTCGCACCTGGATACTCAGCCTACCGGAGGAAAATTTGA
CGGCGTCTTAGGTGTATTGAGCGGACTGGAAGCTGTCCGTACGATGAATGACTTGGGAATCA
AGACCAAACACCCAATTGTCGTTACCAATTGGACTAACGAGGAGGGGGCGCGCTTTGCACCT
GCAATGTTGGCCTCTGGCGTATTTGCAGGCGTACACACCTTAGAATATGCCTACGCCCGTAA
GGACCCAGAAGGCAAATCTTTCGGTGATGAATTAAAGCGTATCGGCTGGTTGGGCGATGAGG
AAGTTGGTGCGCGTAAAATGCACGCATACTTTGAATACCACATCGAGCAAGGCCCAATTCTT
GAGGCGGAGAATAAGCAAATTGGCGTTGTCACTCATTGTCAAGGCCTGTGGTGGCTGGAATT
TACTTTGACAGGTCGCGAGGCTCATACCGGATCAACACCGATGGATATGCGTGTGAATGCGG
GATTAGCGATGGCTCGTATTCTTGAAATGGTCCAAACCGTTGCCATGGAGAACCAACCCGGT
GCAGTGGGCGGCGTGGGGCAGATGTTTTTTAGCCCTAACTCCCGCAACGTGTTACCGGGAAA
AGTAGTTTTCACAGTAGACATCCGTTCCCCTGATCAAGCAAAGCTGGATGGAATGCGCGCAC
GCATTGAGGCAGAAGCGCCAAAAATTTGTGAGCGTCTGGGTGTCGGATGCTCGATTGAAGCA
GTAGGTCACTTCGACCCAGTAACCTTTGACCCTAAACTTGTAGAGACAGTTCGTGGGGCGGC
AGAGAAATTAGGGTATTCGCATATGAACTTAGTTTCTGGCGCCGGACACGACGCCTGCTGGG
CGGCTAAGGTTGCACCTACGACAATGATCATGTGTCCTTGTGTGGGCGGACTTTCGCATAAT
GAGGCGGAAGATATTTCTCGTGAATGGGCAGCAGCCGGTGCAGACGTCCTTTTTCATGCTGT
TTTGGAAACCGCTGAGATTGTGGAGTGACTTGGGAGACCAACTA

#### 5. Supporting Figures

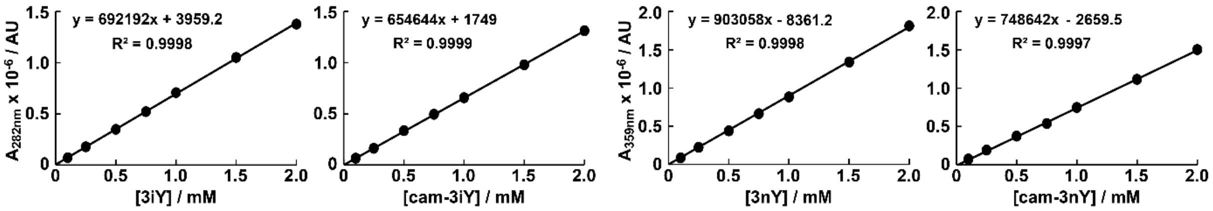


**Figure S1:** HPLC calibration curves for ncAAs and carbamoylated precursors.

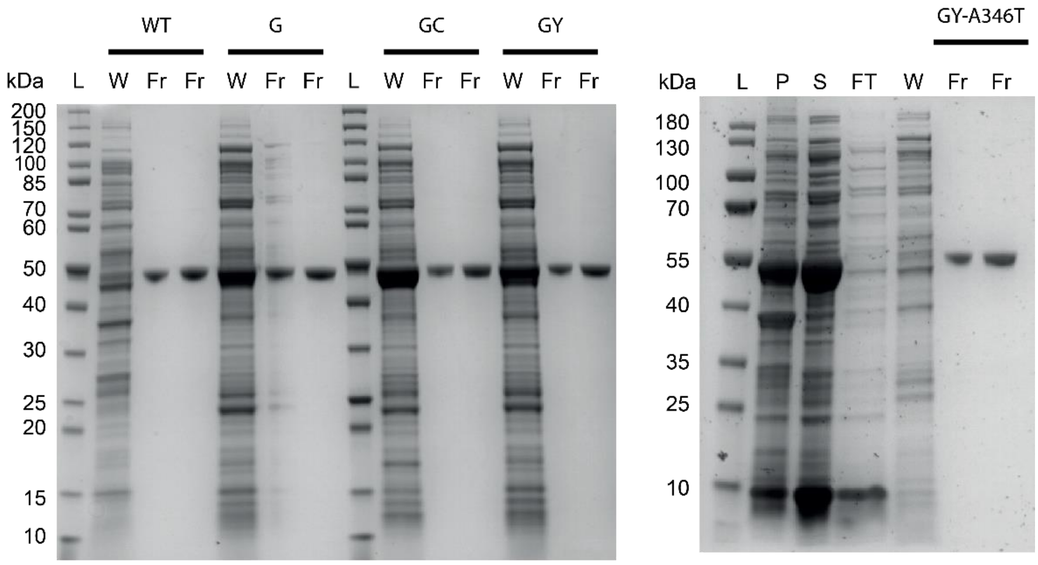


**Figure S2:** SDS-PAGE analysis showing that SmLcar variants (~47 kDa) were obtained in high purity. L = ladder, W = washes, Fr = elution fraction, P = pellet, S = soluble fraction, FT = flowthrough.

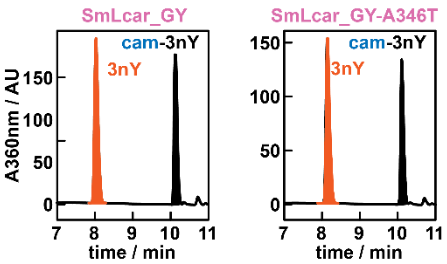


**Figure S3:** HPLC chromatograms showing that the reversion of the Ala346Thr substitution in SmLcar_GY does not drastically alter the activity of the enzyme for cam-3nY.

#### 6. NMR spectra

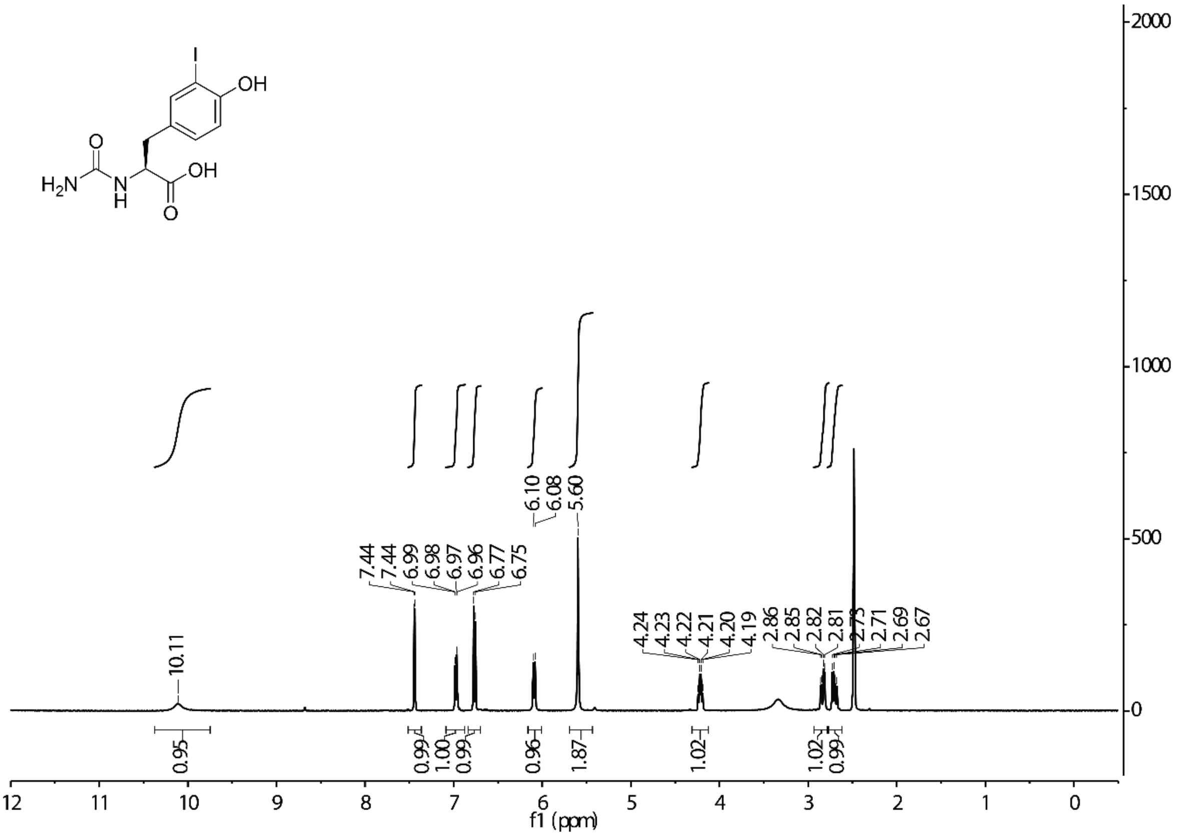


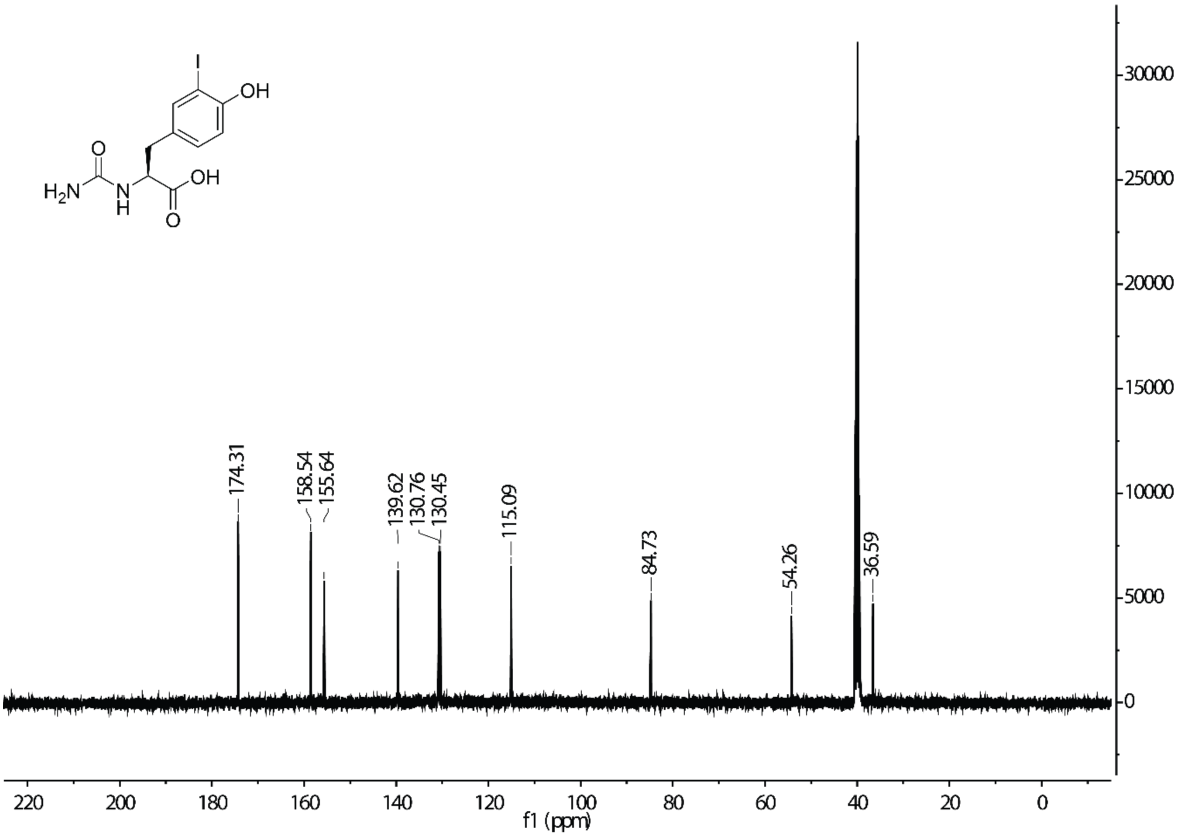

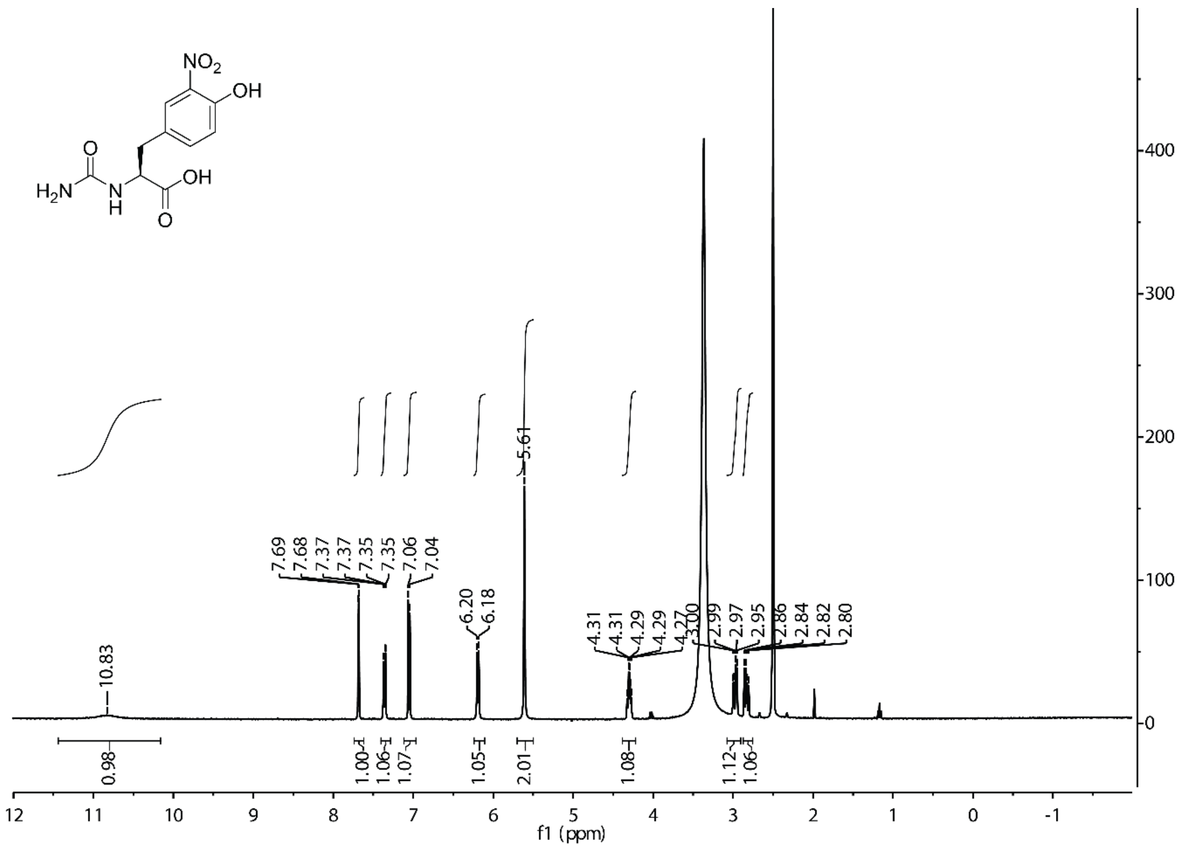

NO2
OH
O
H2N
N
H
OH
O
10.83
7.69
7.68
7.37
7.37
7.35
7.35
7.06
7.04
6.20
6.18
5.61
4.31
4.31
4.29
4.29
4.27
3.00
2.99
2.97
2.95
2.86
2.84
2.82
2.80
0.98
1.00
1.06
1.07
1.05
2.01
1.08
1.12
1.06
f1 (ppm)


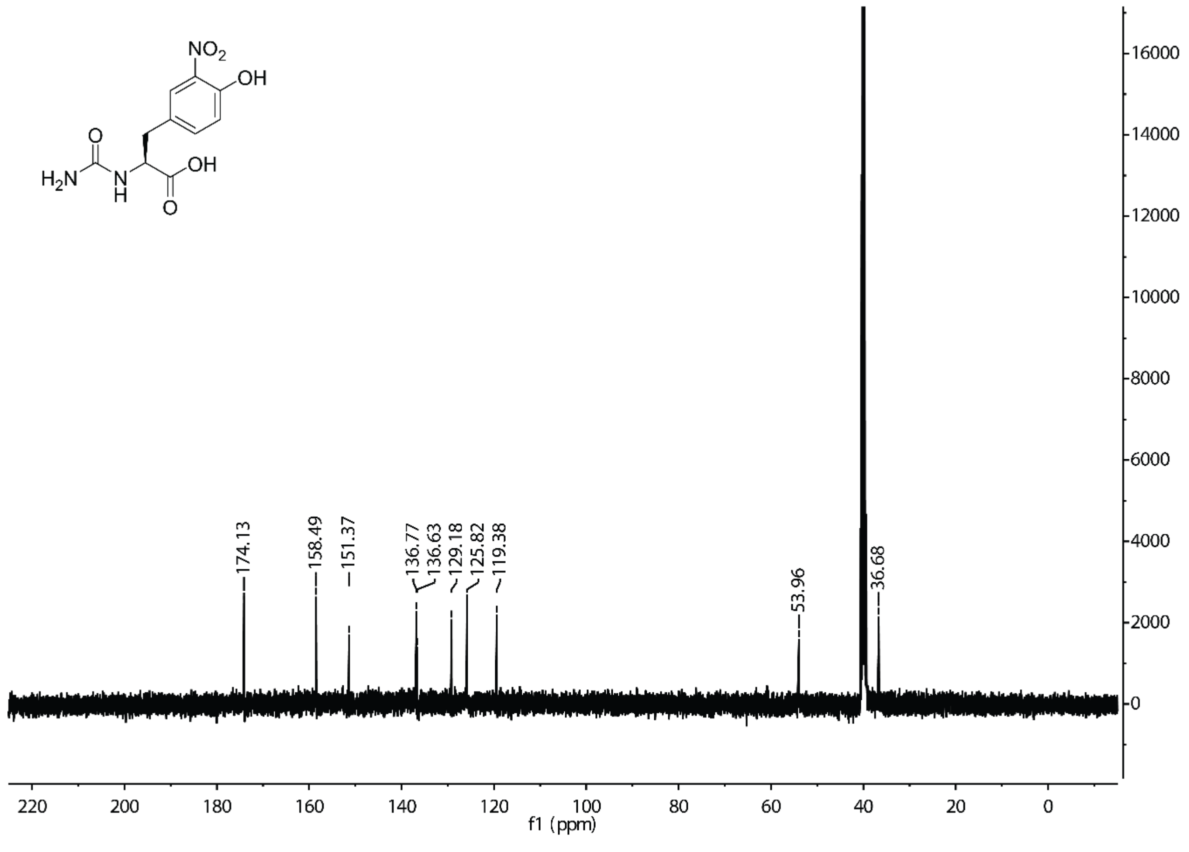

NO2
OH
O
H2N
N
H
OH
O
174.13
158.49
151.37
136.77
136.63
129.18
125.82
119.38
53.96
36.68
f1 (ppm)

### 7. References

1 Shin, I., Han, K. & Rhee, S. Structural Insights into the Substrate Specificity of (S)-Ureidoglycolate Amidohydrolase and Its Comparison with Allantoate Amidohydrolase. *Journal of Molecular Biology* **426**, 3028-3040 (2014).

2 Martínez-Rodríguez, S. *et al.* Mutational and Structural Analysis of <span class="smallcaps smallerCapital">l</span>-<i>N</i>-Carbamoylase Reveals New Insights into a Peptidase M20/M25/M40 Family Member. *Journal of Bacteriology* **194**, 5759-5768 (2012).

3 Martínez-Rodríguez, S., Clemente-Jiménez, J. M., Rodríguez-Vico, F. & Las Heras-Vázquez, F. J. Molecular Cloning and Biochemical Characterization of <i>L</i>-N-Carbamoylase from <i>Sinorhizobium meliloti</i> CECT4114. *Microbial Physiology* **9**, 16-25 (2005).

4 Martínez-Rodríguez, S. *et al.* Thermodynamic and mutational studies of l-N-carbamoylase from Sinorhizobium meliloti CECT 4114 catalytic centre. *Biochimie* **88**, 837-847 (2006).

5 Zheng, H. *et al.* CheckMyMetal: a macromolecular metal-binding validation tool. *Acta Crystallogr D Struct Biol* **73**, 223-233 (2017).

6 Sakamoto, K. *et al.* Genetic encoding of 3-iodo-L-tyrosine in Escherichia coli for single-wavelength anomalous dispersion phasing in protein crystallography. *Structure* **17**, 335-344 (2009).

7 M. Lenman, M., Lewis, A. & Gani, D. Synthesis of fused 1,2,5-triazepine-1,5-diones and some N2- and N3-substituted derivatives: potential conformational mimetics for cis-peptidyl prolinamides 1. *Journal of the Chemical Society, Perkin Transactions 1*, 2297-2312 (1997).

8 Engler, C., Kandzia, R. & Marillonnet, S. A one pot, one step, precision cloning method with high throughput capability. *PLoS One* **3**, e3647 (2008).

9 Tack, D. S. *et al.* Addicting diverse bacteria to a noncanonical amino acid. *Nature Chemical Biology* **12**, 138-140 (2016).

10 Rubini, R. & Mayer, C. Addicting Escherichia coli to New-to-Nature Reactions. *ACS Chem Biol* **15**, 3093-3098 (2020).

11 Mirdita, M. *et al.* ColabFold: making protein folding accessible to all. *Nat Methods* **19**, 679-682 (2022).

12 Martinez-Rodriguez, S., Martinez-Gomez, A. I., Rodriguez-Vico, F., Clemente-Jimenez, J. M. & Las Heras-Vazquez, F. J. Carbamoylases: characteristics and applications in biotechnological processes. *Appl Microbiol Biotechnol* **85**, 441-458 (2010).

13 Sumbalova, L., Stourac, J., Martinek, T., Bednar, D. & Damborsky, J. HotSpot Wizard 3.0: web server for automated design of mutations and smart libraries based on sequence input information. *Nucleic Acids Res* **46**, W356-W362 (2018).

14 Bowler, M. W. *et al.* MASSIF-1: a beamline dedicated to the fully automatic characterization and data collection from crystals of biological macromolecules. *J Synchrotron Radiat* **22**, 1540-1547 (2015).

15 Kabsch, W. Integration, scaling, space-group assignment and post-refinement. *Acta Crystallogr D Biol Crystallogr* **66**, 133-144 (2010).

16 Evans, P. Scaling and assessment of data quality. *Acta Crystallogr D Biol Crystallogr* **62**, 72-82 (2006).

17 Collaborative Computational Project, N. The CCP4 suite: programs for protein crystallography. *Acta Crystallogr D Biol Crystallogr* **50**, 760-763 (1994).

18 McCoy, A. J. Solving structures of protein complexes by molecular replacement with Phaser. *Acta Crystallogr D Biol Crystallogr* **63**, 32-41 (2007).

19 Matthews, B. W. Solvent content of protein crystals. *Journal of Molecular Biology* **33**, 491-497 (1968).

20 Emsley, P. & Cowtan, K. Coot: model-building tools for molecular graphics. *Acta Crystallogr D Biol Crystallogr* **60**, 2126-2132 (2004).

21 Murshudov, G. N. *et al.* REFMAC5 for the refinement of macromolecular crystal structures. *Acta Crystallogr D Biol Crystallogr* **67**, 355-367 (2011).

22 Zheng, H. *et al.* Validation of metal-binding sites in macromolecular structures with the CheckMyMetal web server. *Nat Protoc* **9**, 156-170 (2014).

23 Joosten, R. P., Long, F., Murshudov, G. N. & Perrakis, A. The PDB_REDO server for macromolecular structure model optimization. *IUCrJ* **1**, 213-220 (2014).

24 Chen, V. B. *et al.* MolProbity: all-atom structure validation for macromolecular crystallography. *Acta Crystallogr D Biol Crystallogr* **66**, 12-21 (2010).

25 Tian, W., Chen, C., Lei, X., Zhao, J. & Liang, J. CASTp 3.0: computed atlas of surface topography of proteins. *Nucleic Acids Res* **46**, W363-W367 (2018).

26 Liebschner, D. *et al.* Polder maps: improving OMIT maps by excluding bulk solvent. *Acta Crystallogr D Struct Biol* **73**, 148-157 (2017).
